# Genetic dysregulation of an endothelial Ras signaling network in vein of Galen malformations

**DOI:** 10.1101/2023.03.18.532837

**Authors:** Shujuan Zhao, Kedous Y. Mekbib, Martijn A. van der Ent, Garrett Allington, Andrew Prendergast, Jocelyn E. Chau, Hannah Smith, John Shohfi, Jack Ocken, Daniel Duran, Charuta G. Furey, Hao Thi Le, Phan Q. Duy, Benjamin C. Reeves, Junhui Zhang, Carol Nelson-Williams, Di Chen, Boyang Li, Timothy Nottoli, Suxia Bai, Myron Rolle, Xue Zeng, Weilai Dong, Po-Ying Fu, Yung-Chun Wang, Shrikant Mane, Paulina Piwowarczyk, Katie Pricola Fehnel, Alfred Pokmeng See, Bermans J. Iskandar, Beverly Aagaard-Kienitz, Adam J. Kundishora, Tyrone DeSpenza, Ana B.W. Greenberg, Seblewengel M. Kidanemariam, Andrew T. Hale, James M. Johnston, Eric M. Jackson, Phillip B. Storm, Shih-Shan Lang, William E. Butler, Bob S. Carter, Paul Chapman, Christopher J. Stapleton, Aman B. Patel, Georges Rodesch, Stanislas Smajda, Alejandro Berenstein, Tanyeri Barak, E. Zeynep Erson-Omay, Hongyu Zhao, Andres Moreno-De-Luca, Mark R. Proctor, Edward R. Smith, Darren B. Orbach, Seth L. Alper, Stefania Nicoli, Titus J. Boggon, Richard P. Lifton, Murat Gunel, Philip D. King, Sheng Chih Jin, Kristopher T. Kahle

**Affiliations:** Department of Genetics, School of Medicine, Washington University, St. Louis, MO, USA; Department of Neurosurgery, Massachusetts General Hospital, Harvard Medical School, Boston, MA, USA; Department of Neurosurgery, Yale University School of Medicine, New Haven, CT, USA; Department of Microbiology and Immunology, University of Michigan Medical School, Ann Arbor, MI, USA; Department of Pathology, Yale University School of Medicine, New Haven, CT, USA; Yale Zebrafish Phenotyping Core, Yale University School of Medicine, New Haven, CT, USA; Department of Molecular Biophysics and Biochemistry, Yale University School of Medicine, New Haven, CT, USA; Department of Neurosurgery, University of Mississippi Medical Center, Jackson, MS, USA; Department of Genetics, Yale University School of Medicine, New Haven, CT, USA; Department of Biostatistics, Yale School of Public Health, New Haven, CT, USA; Yale Genome Editing Center, Department of Comparative Medicine, Yale School of Medicine, New Haven, CT, USA; Laboratory of Human Genetics and Genomics, The Rockefeller University, New York, NY, USA; Department of Neurosurgery, Boston Children’s Hospital, Harvard Medical School, Boston, MA, USA; Department of Neurological Surgery, University of Wisconsin School of Medicine and Public Health, Madison, WI, USA; Department of Radiology, University of Wisconsin School of Medicine and Public Health, Madison, WI, USA; Department of Biochemistry, Microbiology and Immunology, University of Ottawa, Ottawa, ON, Canada; Department of Neurosurgery, University of Alabama School of Medicine, Birmingham, AL, USA; Department of Neurosurgery, Johns Hopkins University School of Medicine, Baltimore, MD, USA; Department of Neurosurgery, Hospital of the University of Pennsylvania, Philadelphia, PA, USA; Division of Neurosurgery, Children’s Hospital of Philadelphia, Philadelphia, PA, USA; Service de Neuroradiologie Diagnostique et Thérapeutique, Hôpital Foch, Suresnes, France; Department of Neurosurgery, Icahn School of Medicine at Mount Sinai, New York, NY, USA; Department of Radiology, Autism & Developmental Medicine Institute, Genomic Medicine Institute, Geisinger, Danville, PA, USA; Department of Neurointerventional Radiology, Boston Children’s Hospital, Harvard Medical School, Boston, MA, USA; Division of Nephrology and Center for Vascular Biology Research, Beth Israel Deaconess Medical Center, and Department of Medicine, Harvard Medical School, Boston, MA, USA; Department of Pharmacology, Yale University School of Medicine, New Haven, CT, USA; Yale Cardiovascular Research Center, Department of Internal Medicine, Section of Cardiology, Yale University School of Medicine, New Haven, CT, USA; Department of Pediatrics, Washington University School of Medicine, St. Louis, MO, USA; Division of Genetics and Genomics, Boston Children’s Hospital, Boston, MA, US; Broad Institute of MIT and Harvard, Cambridge, MA, USA

## Abstract

To elucidate the pathogenesis of vein of Galen malformations (VOGMs), the most common and severe congenital brain arteriovenous malformation, we performed an integrated analysis of 310 VOGM proband-family exomes and 336,326 human cerebrovasculature single-cell transcriptomes. We found the Ras suppressor p120 RasGAP (*RASA1*) harbored a genome-wide significant burden of loss-of-function *de novo* variants (p=4.79×10^-7^). Rare, damaging transmitted variants were enriched in Ephrin receptor-B4 (*EPHB4*) (p=1.22×10^-5^), which cooperates with p120 RasGAP to limit Ras activation. Other probands had pathogenic variants in *ACVRL1*, *NOTCH1*, *ITGB1*, and *PTPN11*. *ACVRL1* variants were also identified in a multi-generational VOGM pedigree. Integrative genomics defined developing endothelial cells as a key spatio-temporal locus of VOGM pathophysiology. Mice expressing a VOGM-specific *EPHB4* kinase-domain missense variant exhibited constitutive endothelial Ras/ERK/MAPK activation and impaired hierarchical development of angiogenesis-regulated arterial-capillary-venous networks, but only when carrying a “second-hit” allele. These results illuminate human arterio-venous development and VOGM pathobiology and have clinical implications.

## INTRODUCTION

The development of the cerebrovascular system, which is required to meet the hemodynamic and nutritive demands of embryogenesis, is a complex yet reproducible process dictated by spatially and temporally coordinated events that are regulated by genetically determined factors ^1, 2^. These factors control changes in gene expression that regulate vasculogenesis, angiogenesis, and arterio-venous specification ^3–5^, which give rise to distinct, contiguous cerebral vessels with segments identified as arteries, capillaries, and veins ^2^. The heterogeneous cell composition along this hierarchically-organized arterio-venous axis includes endothelial cells, pericytes, smooth muscle cells, and other perivascular cells (e.g., neurons and immune cells) ^1, 6, 7^. Although experiments in model systems, including chick embryos, zebrafish, and mice, have detailed numerous coordinated molecular interactions within and among these cells that endow the cerebrovasculature with its specialized structural and functional properties, the genetic regulation of arteriovenous development in humans remains poorly understood ^2^. The genomic study of rare, severe congenital cerebrovascular anomalies can help elucidate the genes and pathways that are essential for human cerebrovasculature development and identify potential therapeutic targets relevant for more common forms of disease ^8, 9^.

During normal brain development, primitive choroidal and subependymal arteries that perfuse deep brain structures are connected via a primitive meningeal capillary network to the embryonic precursor of the vein of Galen termed the median prosencephalic vein of Markowski (MPV). The MPV returns deep cerebral venous blood to dural sinuses that drain into the internal jugular veins ^10^. Vein of Galen malformations (VOGMs), the most common and severe arteriovenous malformations (AVMs) of the human neonatal brain ^11, 12^, directly connect primitive choroidal or subependymal cerebral arteries to the MPV without an intervening capillary network. This pathological connection exposes the MPV to dangerously high blood flow and blood pressures that can lead to high-output cardiac failure, hydrocephalus associated with venous congestion, and intracerebral hemorrhage ^13^ (**Fig. 1a**). VOGMs can also be associated with other neurodevelopmental pathology and structural heart defects ^14^, and can be fatal if untreated. Although VOGM treatment has greatly benefited from recent advances in endovascular therapy ^15^, the morbidity and mortality related to VOGM remain high, even at specialized referral centers ^16, 17^.

**Figure 1.**
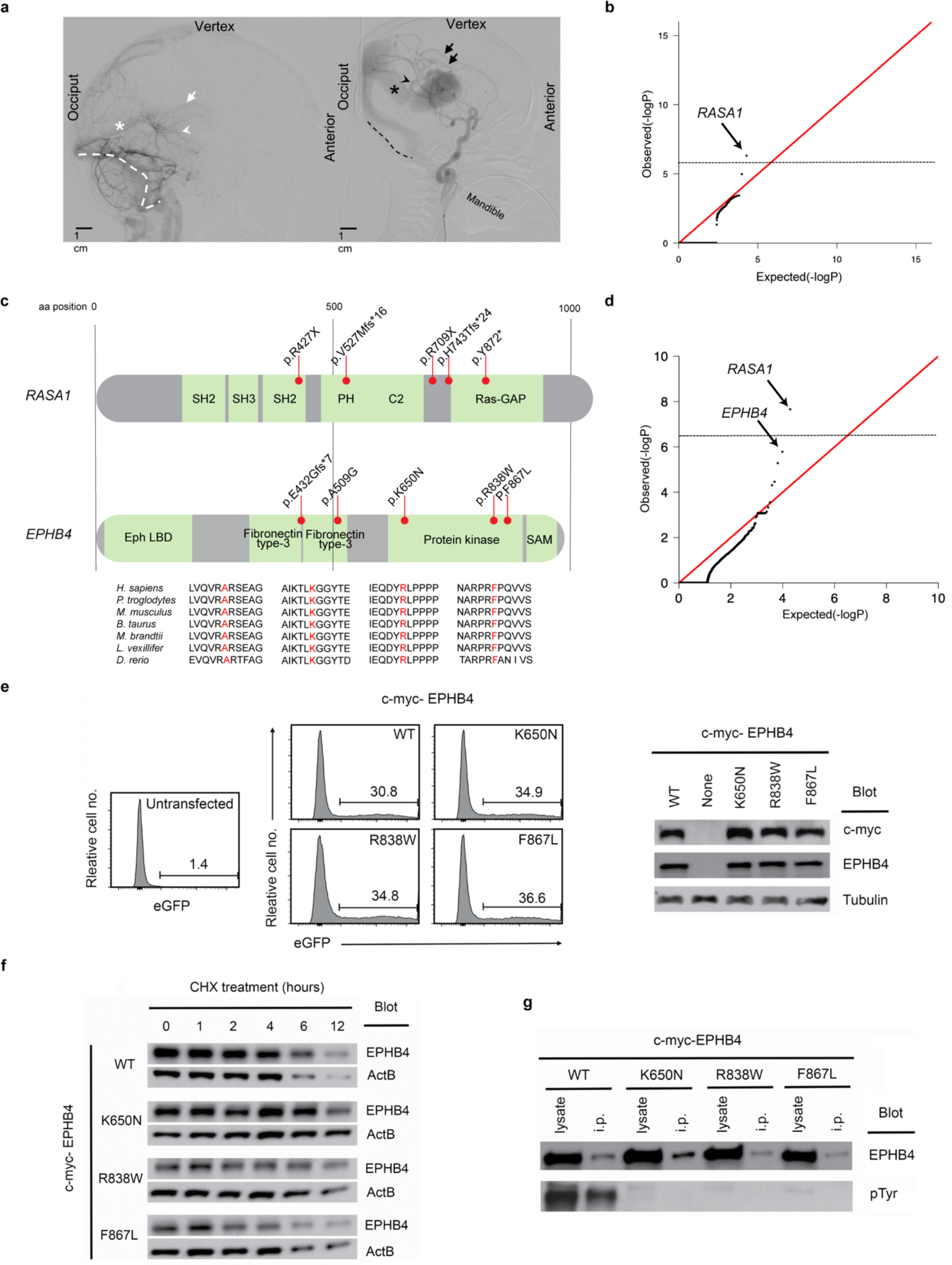
VOGM-associated variants in *RASA1* and *EPHB4*. (a) Representative digital subtraction angiography images of normal cerebrovascular anatomy (left) and VOGM anatomy (right). In normal cerebrovascular anatomy, the deep veins drain into the straight sinus (white asterisk), whereas in VOGM cerebrovascular anatomy the vein of Galen most commonly drains into a persistent falcine sinus (black asterisk). The normal deep venous anatomy visualized includes the internal cerebral veins (white arrow) and the basal vein of Rosenthal (white arrowhead). In contrast, the VOGM image shows an arteriovenous shunt with arteries visible concurrently with veins and venous sinuses. This malformation involves prominent splenial vessels deriving from the anterior cerebral artery (black arrows) and posterior choroidal arteries (black arrowhead). Other cases can involve more prominent thalamoperforators as well. The normal drainage of the torcula typically involves the bilateral transverse and sigmoid sinuses (white dashed line). Additionally, in abnormal venous anatomy, VOGMs typically recruit dilated occipital sinuses (black dashed line). (b) Quantile-quantile plot of observed versus expected P value for rare damaging (D-mis + LoF) DNVs with MAF σ:4 x 10^-4^ in the Exome Aggregation Consortium database for all genes. *RASA1* exhibits exome-wide significant enrichment for all rare variants in VOGM cases. The exome-wide significant cutoff was 8.6 × 10^−7^ (0.05 / (3 × 19,347)). D-mis are missense mutations predicted to be deleterious per MetaSVM or MPC σ: 2. LoF mutations include stop-gain, stop-loss, frameshift insertions/deletions, canonical splice sites, and start-loss. LoF and D-mis mutations were considered ‘damaging’. DNV, *de novo* variant. MAF, minor allele frequency. (c) RASA1 and EPHB4 functional domains (green rectangles) with the location of VOGM mutations and phylogenetic conservation of wild-type amino acid (red text) at each mutated position. 5 LoF mutations were found in *RASA1*, including 2 DNVs (SH2 domain stop-gain variant p. R427X, PH domain frameshift variant p. V527Mfs*16), and 3 transmitted variants (frameshift p. H743fs*24, stop-gain p. R709X and Ras-GAP domain stop-gain variant p. Y872*). Transmitted damaging mutations in *EPHB4* including 2 variants that are in the Fibronectin type III domain (frameshift variant p. E432Gfs*7 and D-mis variant p. A509G), and 3 D-mis variants (p. K650N, p. R838W, p. F867L) located in the Protein kinase domain. The D-mis variants in EPHB4 are in regions that are highly conserved among different species. SH2, Src homology 2. PH, Pleckstrin Homology. (d) Quantile-quantile plots of observed versus expected P value for rare damaging (D-mis + LoF) variants with MAF σ: 5 x 10^-5^ in the Genome Aggregation database (gnomAD) from case-control burden test. The genome-wide significant cutoff was 2.6 × 10^−6^ (0.05 of 19,347). (e) Steady-state abundance of EPHB4 D-mis mutants. Cos-7 cells were transiently transfected with c-myc-tagged WT or mutant EPHB4 as indicated. Cells were co-transfected with an eGFP-encoding vector to assess comparable transfection efficiency determined by flow cytometry. Percentage of eGFP+ cells with EPHB4 D-mis mutants is shown. EPHB4 abundance was determined in cell lysates by EPHB4 and c-myc immunoblot. Blots were probed for tubulin to demonstrate equivalent protein loading. Abundance of Lys650Asn, Arg838Trp, and Phe867Leu EPHB4 was each comparable to that of WT EPHB4 when expressed in Cos-7 cells. (f) Stability of EPHB4 D-mis mutants. Cells were treated with cycloheximide (CHX) for the indicated times before lysis, and EPHB4 abundance was determined by immunoblot. Blots were probed for actin to compare EPHB4 decay rates to that of an endogenous cell reference protein. After cycloheximide treatment to block protein translation, the rate of decay of EPHB4 mutants was similar to that of WT EPHB4. (g) Phosphotyrosine content of VOGM-associated EPHB4 kinase domain mutants. Phosphotyrosine content (pTyr) was determined by immunoblot of whole cell lysates and EPHB4 immunoprecipitates.

Our limited understanding of the molecular pathophysiology of VOGMs has hindered the development of early diagnostic tests and non-procedural therapeutic strategies. Early hypotheses proposed VOGMs were secondary to thrombosis of the straight sinus ^18–21^. However, newer theories suggest VOGMs may instead reflect failed angiogenic intussusception during the development of the choroid plexuses, with ischemia playing a potential role ^22^. Although most VOGMs are sporadic lesions, VOGMs have also been associated with several Mendelian disorders, sometimes co-existing with other intracranial and extracranial AVMs and multifocal capillary malformations and telangiectasias ^23–26^. These include several cases of autosomal dominant (AD) capillary malformation-AVM syndrome types 1 and 2 (CM-AVM1/2), caused by *RASA1* mutation (OMIM: 608354) ^27^ or *EPHB4* mutation (OMIM: 618196) ^25^, and single VOGM cases in AD hereditary hemorrhagic telangiectasia type 1 and 2 (HHT1/2) caused by mutation in *ENG* (OMIM: 187300) ^28^ or *ACVRL1* (OMIM: 600376) ^29^. These data implicate genetic contributions to VOGM, but the underlying genetic cause of most cases remains unknown. Moreover, the cellular and molecular mechanism of VOGM-associated mutations are poorly understood due in part to a lack of adequate mammalian models.

The rare and overwhelmingly sporadic nature of VOGM cases ^30–32^ has limited the power of traditional human genetic approaches (including linkage and genome-wide association studies) to identify causative genes for VOGM and other congenital cerebrovascular lesions. These limitations motivate the curation of deeply-phenotyped proband-parent (“trio”) based cohorts, recruited through multi-institutional, international collaboration. Whole-exome sequencing (WES) and computational statistical analyses can then be applied to search for rare damaging mutations in probands more often than expected by chance ^26^. This agnostic genomic approach has aided gene discovery for other brain and cerebrovascular malformations ^33–36^, congenital heart disease ^37^, congenital hydrocephalus ^38, 39^, and other genetically heterogeneous neurodevelopmental disorders including autism, craniosynostosis, and epilepsy ^40–45^. In addition, recent single-cell RNA-sequencing (scRNA-seq) studies have begun to systematically catalog cell types of the developing and mature human cerebrovasculature, expanding knowledge about perivascular cell diversity and defining endothelial cell molecular signatures correlated with arteriovenous segmentation ^46, 47^. Integration of WES findings with other large-scale-omic datasets can help elucidate the cellular and molecular mechanisms of disease genes by defining their spatio-temporal expression and associated transcriptional and protein-protein interaction networks ^36, 39, 48^.

Here, we aimed to integrate trio-based WES data of the largest VOGM cohort collected to date (310 VOGM proband-family exomes) with 336,326 human cerebrovasculature single-cell transcriptomes in an integrated, systems-level investigation of VOGM pathogenesis (see **Graphical Abstract**). We hypothesized that: (i) multiple novel VOGM candidate genes harboring pathogenic *de novo* and transmitted variants will be discovered using trio-based WES; (ii) VOGM genes will spatiotemporally converge in co-expression modules, cell types, and biological pathways pertinent to the regulation of endothelial biology in ways distinct from those of other congenital cerebrovascular diseases; (iii) systematic comparison of phenotypic data from individual VOGM cases will assist gene discovery by clustering cases with similar endophenotypes, thereby defining clinically-relevant disease subclasses; and (iv) functional studies of candidate variants in model systems can increase the confidence of gene pathogenicity and provide insight into mutation mechanisms.

## RESULTS

### WES of the largest trio-based VOGM cohort to date

We ascertained a total of 114 probands with radiographically confirmed VOGM (see **Methods**) treated by endovascular therapy, including 90 proband-parent trios (each with a single affected offspring), 13 duo cases, and 11 singleton cases (**Table S1**). These included 55 previously described VOGM probands ^26^. Among 114 VOGM probands, 34.2% were female; 78.9% were self-reported Europeans; 60% of probands were diagnosed prenatally or within one month after birth; only 3.8% were diagnosed after age 2. Salient features at diagnosis included developmental delay (54%), macrocephaly (48%), hydrocephalus (48%), prominent face and/or scalp vasculature (45%), cutaneous vascular lesions (22%), and congestive heart failure (40%). Also present were structural heart defects (7%), including partial anomalous pulmonary venous return, patent ductus arteriosus, and pulmonary valve stenosis. **Table S2** summarizes the demographics and clinical features of our VOGM cohort. **Fig. S1** depicts representative imaging for VOGM probands.

DNA was isolated, and WES was performed as previously described ^26^. In parallel, WES of 1,798 control trios comprising parents and unaffected siblings of autism probands were analyzed ^49, 50^ by our in-house informatics pipeline. 92.7% or more of targeted bases had eight or more independent reads in both cases and controls, and 87.8% or more had 15 or more independent reads (see **Table S3** for exome sequence metrics). Variant calling was performed utilizing a combination of the Genome Analysis Toolkit (GATK) HaplotypeCaller ^51, 52^ and Freebayes ^53^. Population allele frequencies were annotated by the Genome Aggregation Database (gnomAD v.2.1.1) and the BRAVO databases ^54^. *De novo* variant (DNV) identification was performed by TrioDeNovo ^55^. MetaSVM and MPC algorithms were used to infer the effect of missense mutations ^56, 57^. Missense variants were considered damaging (D-mis) when predicted to be deleterious by MetaSVM, or if they had an MPC-score ≥ 2. Inferred loss-of-function (LoF) mutations, including stop-gains, stop-losses, frameshift insertions, deletions, and canonical splice-site mutations were considered damaging. Mutations in genes of interest were validated by PCR amplification and Sanger sequencing (**Fig. S2 and S3**).

### Variants in CM-AVM genes *RASA1* and *EPHB4*

We first investigated the contribution of DNVs to VOGM pathogenesis (**Table S4**). The average DNV rate of 1.19 per subject (**Table 1** and **Table S5**) resembled previous results obtained from a similar sequencing platform ^39^ and followed a Poisson distribution (**Fig. S4)**. The burden of DNVs in the control cohort was comparable (**Table S5**). There was no enrichment of synonymous or missense DNVs inferred to be tolerated in VOGM cases. In contrast, D-mis (1.65-fold, P =1.86×10^-2^) and protein-damaging DNVs (LoF + D-mis) (1.50-fold, P =1.72×10^-2^) were enriched among all genes in cases but not in controls (**Table 1**). *RASA1* (probability of loss-of-function intolerance [pLI] = 1.00 in gnomAD v.2.1.1) was the single gene to harbor a genome-wide significant burden of damaging DNVs (one-tailed Poisson p = 4.79×10^-7^) with two novel LoF DNVs (**Fig. 1B and Table S6**). The only other gene with more than one protein-altering DNV (p.Gly202Ser and p.Gln321*) was endothelin-3-converting enzyme *ECE3* (*KEL*) (one-tailed Poisson p = 1.03×10^-5^). From the observed fraction of patients with damaging DNVs we infer that damaging DNVs account for >12% of VOGM cases (**Table 1** and see **Methods**).

**Table 1.**
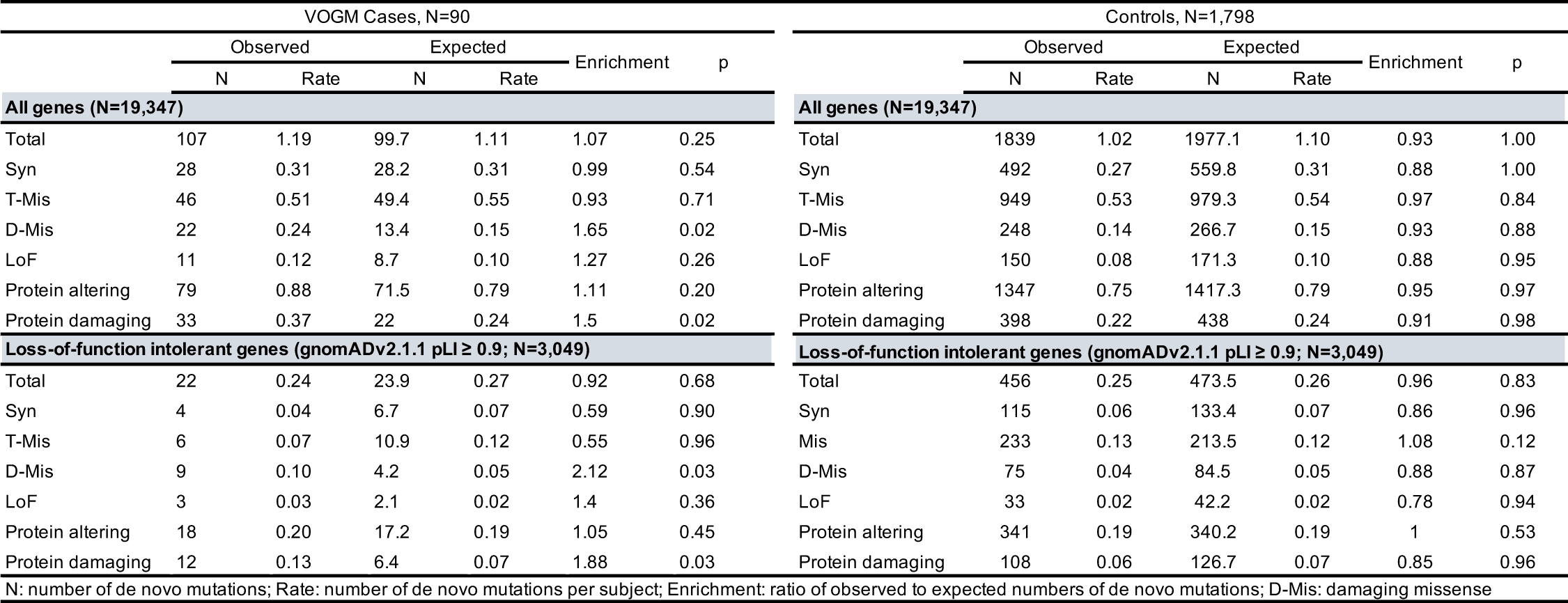
De novo variant enrichment analysis for each functional class in 90 VOGM trio cases and control cohorts

*RASA1*, encoding the negative Ras regulator p120 RasGAP ^58^, contained two novel LoF DNVs, p.Arg427* and p.Val527Mfs*16, in unrelated VOGM probands KVOGM71-1 and KVOGM122-1, respectively (**Table 2 and Fig. S2**). Variants in *RASA1* have been implicated in autosomal dominant type 1 capillary malformation-arteriovenous malformation (CM-AVM1) (OMIM# 608354) ^59^, featuring fast-flow vascular malformations, including systemic and intracranial arteriovenous fistulas and AVMs and, rarely, VOGMs ^27, 30, 60^. Both KVOGM71-1 and KVOGM122-1 had cutaneous capillary malformations characteristic of CM-AVM type 1. See **Fig. S5** for available cutaneous manifestations in VOGM probands and their family members.

**Table 2.** Characteristics of Patients With Variants in RASA1 and EPHB4

| Proband ID | Ethnicity | Sex | Type | Proband Phenotype | Carrier Parent Phenotype | Class | Gene | Position (GRCH37) | AA Change | Bravo WGS Freq | gnomAD WES Freq |
| --- | --- | --- | --- | --- | --- | --- | --- | --- | --- | --- | --- |
| KVOGM_71-1 | African | Female | DNM | Cutaneous vascular lesion | NA | stopgain | *RASA1* | 5:86648999:C:T | p.R427X | NA | NA |
| KVOGM122-1 | European | Male | DNM | Cutaneous vascular lesion | NA | frameshift | *RASA1* | 5:86659286:TTCTG:T | p.V527Mfs*16 | NA | NA |
| KVOGM95-1 | European | Female | Transmitted from mother | Deaf left ear | Cutaneous vascular lesions (mother) | frameshift | *RASA1* | 5:86672737:T:TC | p.H743Tfs*24 | NA | NA |
| KVOGM42-1* | Mexican | Female | Transmitted from mother | Cutaneous vascular lesion, cerebral palsy, neurodevelopmental delay, seizures | Cutaneous vascular lesions (mother) | stopgain | *RASA1* | 5:86672323:C:T | p.R709X | NA | NA |
| KVOGM48-1 | European | Female | Transmitted from mother | Cutaneous vascular lesion, neurodevelopmental delay | Cutaneous vascular lesions (mother) | stopgain | *RASA1* | 5:86676336:T:TA | p.Y695* | NA | NA |
| KVOGM18-1* | European | Male | Transmitted from father | Cerebral palsy, hemiplegia, macular scarring, visual neglect, neurodevelopmental delay, seizures | NA | D-mis | *EPHB4* | 7:100403202:A:G | p.F867L | NA | NA |
| KVOGM33-1* | Mexican | Female | Transmitted from mother | Neurodevelopmental delay | Cutaneous vascular lesions (mother) | D-mis | *EPHB4* | 7:100410537:C:A | p.K650N | NA | NA |
| KVOGM25-1* | European | Male | Transmitted from father | Sternum excavatum, neurodevelopmental delay | NA | D-mis | *EPHB4* | 7:100414876:G:C | p.A509G | 2.39E-05 | 1.48E-05 |
| KVOGM115-1* | European | Male | Transmitted from father | Neurodevelopmental delay, seizures | Cutaneous vascular lesions (father) | frameshift | *EPHB4* | 7:100417179:C:CTC | p.E432Gfs*7 | NA | NA |
| KVOGM_72-1 | East Asian | Male | Transmitted from father | NA | NA | D-mis | *EPHB4* | 7:100403289:G:A | p.R838W | NA | 1.39E-05 |

\* These patients have been reported in our previous study (Duran et al. 2019). Abbreviations: AA, amino acid; Bravo WGS Freq, Bravo whole-genome sequencing frequency; gnomAD WES Freq, gnomAD whole-genome sequencing frequency; D-Mis, deleteriousness of missense variants; DNV, de novo variant; gnomAD WES Freq, Gnome aggregation database whole exome sequence frequency. This global frequency of the Bravo database is based on 132 345 genomes and is independent of ethnicity; Gnome Aggregation Database Whole Exome Sequence Frequency. This frequency is based on 125 748 exomes in the gnomAD database (v2.1.1) and is independent of ethnicity.

To identify additional haploinsufficient genes associated with VOGM undetected by DNV analysis, we next assessed the total burden in all probands of rare (Bravo minor allele frequency [MAF] ý 5×10^-5^) *de novo* and transmitted D-mis and LoF variants. The probability of the observed number of rare variants in each gene occurring by chance was calculated by comparing the observed with the expected burden, adjusting for gene mutability as previously described ^39^. Analysis of damaging variants in all genes revealed a genome-wide significant enrichment (Bonferroni multiple testing threshold = 2.6×10^-6^) of variants in *RASA1*, with five total rare LoF variants (one-tailed binomial p = 1.20×10^-6^; enrichment = 28.4-fold; **Table S7**). In addition to the two *de novo* LoF variants identified in *RASA1* (see above), we identified two transmitted stop-gain variants in *RASA1* (p.Arg709* and p.Tyr695*) in unrelated probands. We also found an unphased frameshift variant in *RASA1* (p.His743Thrfs*24) (**Table 2** and **Fig. 1c**). Case-control burden analysis for rare damaging mutations in all probands versus gnomAD controls showed *RASA1* as having a significant mutational burden in VOGM probands (one-tailed Fisher’s p = 2.20×10^-8^, odds ratio = 67.50, 95% confidence interval [CI] [25.76, infinite [Inf]]) (**Fig. 1D** and **Table S8**).

RAS proteins cycle between an active guanosine-triphosphate (GTP)-bound form and an inactive, guanosine-diphosphate (GDP)-bound form, as do other guanine nucleotide-binding proteins (including heterotrimeric G proteins). The weak intrinsic GTPase activity of RAS proteins is greatly enhanced by GTPase-activating proteins (GAPs) such as p120 RasGAP ^61, 62^. Similar to other LoF *RASA1* mutations ^63^, mutations encoding the prematurely terminated mutants p.Arg427*, p.Val527Metfs*16, p.Arg709*, p.Tyr695*, and p.His743Thrfs*24 are all predicted to cause nonsense-mediated mRNA decay and consequently increase downstream Ras/ERK/MAPK signaling activity ^64, 65^.

We also identified enrichment in rare, damaging variants in *EPHB4*, encoding Ephrin receptor B4 (EphB4), which physically interacts and cooperates with p120 RasGAP to limit Ras activation ^66^ (one-tailed binomial p = 1.22×10^-5^; enrichment = 17.5-fold; **Table S7**). Case-control gene burden analyses for damaging variants in all probands showed a significant mutation burden in *EPHB4* versus gnomAD controls (one-tailed Fisher’s p =1.65×10^-6^, odds ratio = 27.39, 95% CI [10.60, Inf]; **Fig. 1d** and **Table S8)**. *EPHB4* variants ^25^ have been reported in type 2 AD Capillary malformation-arteriovenous malformation (CM-AVM2) (OMIM# 618196). In contrast to *RASA1*, all but one of the *EPHB4* variants were D-mis and transmitted; three of these are not observed in ExAC and gnomAD, and two have a MAF of less than 1.48×10^-5^ in gnomAD (**Table 2**).

The only detected LoF variant in *EPHB4* (p.Glu432Glyfs*7) immediately precedes the fibronectin type 3 (fn3) domain and is expected to truncate the protein kinase domain (**Fig. 1C**), and is predicted to result in nonsense-mediated mRNA decay ^67^. All D-mis variants in *EPHB4* alter highly conserved amino acid residues (**Fig. 1c**). p.Lys650Asn and p.Phe867Leu localize to the Eph-B4 tyrosine kinase domain ^68, 69^. p.Lys650Asn is a surface-exposed residue in the β3-αC loop of the EPHB4 tyrosine kinase N-lobe and is located proximal to the putative binding site of the autoinhibitory juxtamembrane region. p.Phe867Leu is a highly conserved, hydrophobic-core residue in the C-lobe of the EPHB4 tyrosine kinase domain. p.Ala509Gly lies in a highly conserved, hydrophobic-core residue in the second of two EPHB4 extracellular fibronectin III domains (**Fig. S2)**. *In silico* biophysical modeling suggests these VOGM-associated D-mis variants significantly disrupt EPHB4 structure and function ^26^. The steady-state abundance of Lys650Asn, Arg838Trp, and Phe867Leu EPHB4 variants resembled that of WT EPHB4 when expressed in Cos-7 cells (**Fig. 1e**). Cycloheximide block of protein translation revealed polypeptide decay rates of EPHB4 VOGM mutants similar to that of WT EPHB4 (**Fig. 1f**), suggesting that none of the *EPHB4* kinase domain mutations appear to affect EPHB4 protein stability. In contrast, the phosphotyrosine content in each of the three VOGM-associated EPHB4 kinase domain mutants were significantly reduced as detected by anti-pTyr immunoblotting in whole-cell lysates and anti-EPHB4 immunoprecipitates (**Fig. 1g**). The findings indicate that VOGM-associated EPHB4 D-mis variants inhibit the protein tyrosine kinase activity of EPHB4.

Patient phenotypes associated with *RASA1* and *EPHB4* variants are described in **Table S9**. These kindreds notably include eight additional family members without diagnosed VOGM who carry the same variants. Among these, three carriers of *RASA1* variants and two carriers of *EPHB4* variants had cutaneous vascular lesions (**Table S9**). In contrast, cutaneous vascular lesions and cardiac abnormalities were absent among confirmed non-carrier family members. For example, KVOGM42-1 with the rare transmitted *RASA1* p.Arg709* variant had a carrier mother without a VOGM, but with cutaneous vascular lesions and an extensive family history on the maternal side of aneurysms, stroke, and Raynaud’s syndrome in several other family members unavailable for sequencing. Similarly, the father of KVOGM115-1 carrying the *EPHB4* p.Glu432Glyfs*7 variant lacked VOGM but had a capillary malformation. The proband also had a brother with multiple cutaneous capillary malformations and other vascular lesions on his face and leg unavailable for sequencing.

Interestingly, before inclusion in this study, none of the VOGM probands or their family members carried a clinical or genetic diagnosis of CM-AVM. Together, these findings show incomplete penetrance and variable expressivity of transmitted variants in the related CM-AVM genes *RASA1* and *EPHB4* in VOGM.

### Variants in *ACVRL1* and other mutation-intolerant Mendelian vascular disease genes

To identify other potential VOGM genes and gain insight into the molecular pathways impacted by their mutation, we performed Gene Ontology (GO), Reactome, and Wiki pathway analyses (see **Methods**) on genes with a pLI > 0.9 that harbored damaging *de novo* and/or rare (MAF ≤ 2 × 10^−5^) damaging transmitted mutations (see **Table S10** and **Methods**). Among the top significantly enriched terms (**Fig. 2a and Fig. S6**) was Reactome pathway term REAC:R-hsa-422475 associated with axon guidance (p=2.17×10^-15^), known to be critical for vascular patterning and regulated by Ephrin-Eph receptor signaling^1^. Other GO biological process and molecular function terms enriched in VOGM probands included those associated with heart development (GO:0007507, p=5.19×10^-10^) and Ras GTPase activator activity (GO:0005096, p=3.35×10^-10^) (**Fig. 2a**).

**Figure 2.**
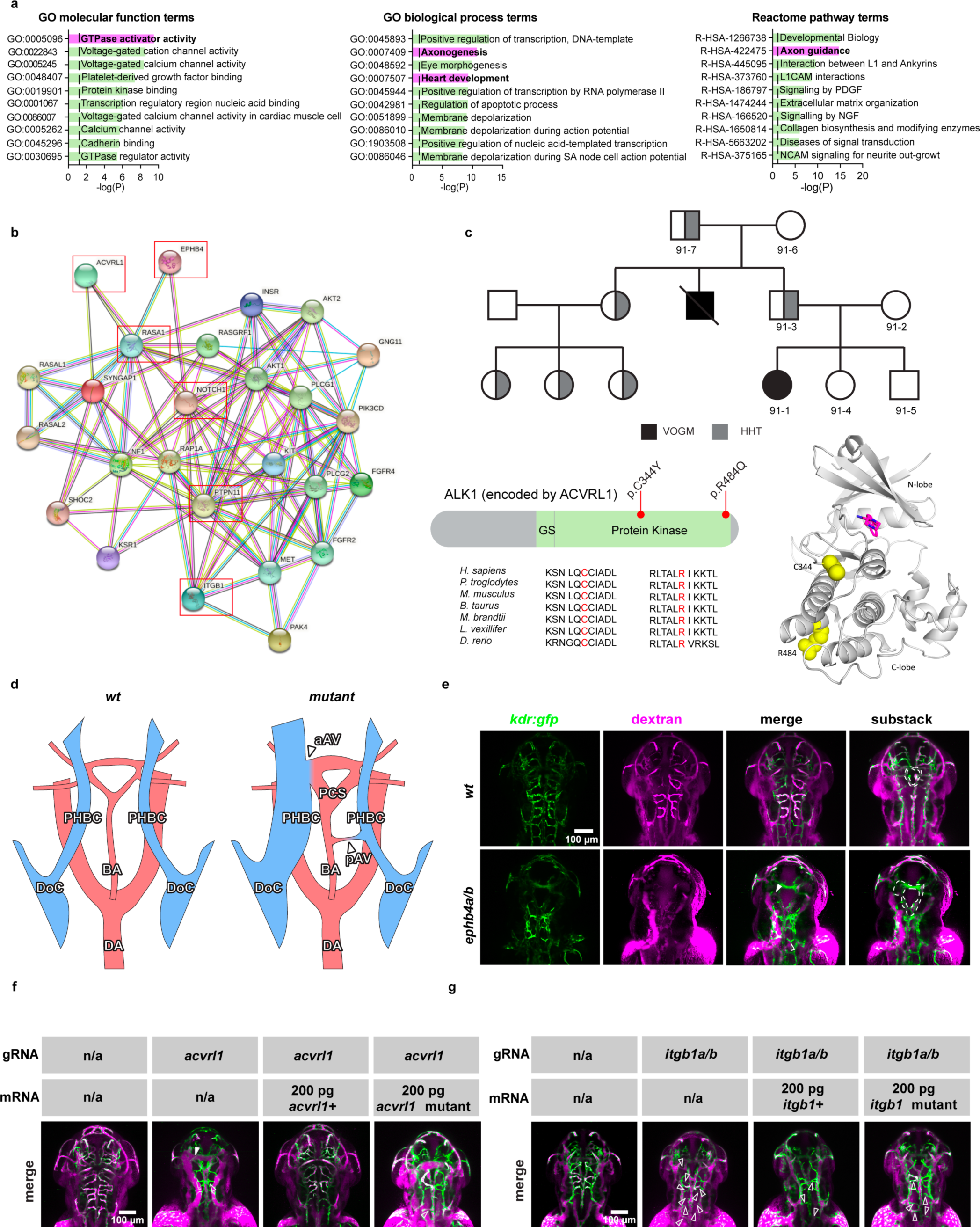
VOGM-associated variants in *ACVRL1* and other Mendelian vascular disease genes. (a) Top 10 GO molecular function, GO biological process, and Reactome pathway enrichment terms. The y-axis depicts GO term and Reactome pathway ID numbers. The x-axis depicts-log (P value) and the dotted line represents the α = 0.05 significance threshold. The GO term and Reactome pathway term name overline their respective bars. (b) Interactome of mutated Ras signaling genes in VOGM. Genes with damaging Ras signaling variants (WP4223 in Wiki pathway analysis) and *EPHB4*, *NOTCH1*, *ACVRL1,* and *ITGB1* (a total of 27 genes) were inputted into String (https://string-db.org/cgi/about.pl) and mapped onto a single STRING interactome. This network has significantly more interactions than expected (PPI enrichment p-value < 1.0×10^-16^). All 6 VOGM risk genes (highlighted by red box) contribute significantly to the PPI enrichment of this network. (c) Multi-generational VOGM family in KVOGM-91. Two transmitted D-mis variants (p. Cys344Tyr and p. Arg484Gln) are in the protein kinase domain. Schematic of the crystal structure of the ALK1 (*ACVRL1*) kinase domain is shown (PDB ID: 3MY0) ^160^. Locations of Cys344 and Arg484 are indicated with yellow spheres. A small molecule inhibitor is bound in the catalytic cleft and colored purple. HHT, hereditary hemorrhagic telangiectasia. (d) Schematic showing layout of cranial arteries (red) and veins (blue) in 48 hpf zebrafish *wt* and mutant embryos. We attempted to identify both anterior abnormal arterial/venous connections (aAV) and posterior abnormal arterial/venous connections (pAV) between the primordial hindbrain channel (PHBC) and basilar artery (BA) or posterior communicating segment (PCS). Also pictured: duct of Cuvier (DoC) and dorsal aorta (DA). Generally, we observed enlarged PHBC and PCS vessels and supernumerary AV connections as mutant phenotypes. (e) Cranial vasculature indicated by Tg(kdr:gfp) and dextran microangiography of 48 hpf wt and ephb4a/b-depleted zebrafish embryo. Scale bar: 100 µm. Cranial vasculature is indicated by Tg(kdr:gfp), dextran microangiogram, and merged channels of the wt and ephb4a/b-deleted embryo. Ventrally restricted confocal substacks emphasizing PCS (dashed lines) are shown. Note both anterior flow-carrying connections between PCS and PHBC (filled arrowheads) and posterior flow-carrying connections between BA and PHBC (open arrowhead). Ventrally restricted confocal substack shows enlarged PCS. (f) Cranial vasculature indicated by Tg(kdr:gfp) and dextran microangiogram of 48 hpf wt and acvrl1-depleted, rescue (with acvrl1+ mRNA) and false rescue (with acvrl1 mutant mRNA) zebrafish embryos. Scale bar: 100 µm. Note massively enlarged PHBC and PCS, and again both anterior flow-carrying connections between PCS and PHBC (filled arrowhead) and posterior flow-carrying connections between BA and PHBC (open arrowhead). In the rescue embryo, PHBC and PCS size has returned to normal with 200 pg acvrl1+. In 48 hpf acvrl1-depleted embryo co-injected with 200 pg acvrl1 Cyc344Tyr mRNA, PHBC and PCS are enlarged, indicating a failure to rescue. (g) Cranial vasculature indicated by Tg(kdr:gfp) and dextran microangiography of 48 hpf wt and itgb1a/b-depleted, rescue (with itgb1+ mRNA) and false rescue (with itgb1 mutant mRNA) zebrafish embryos. Note prevalence of pAVs (open arrowheads) in *itgb1a/b*-depleted embryo. Fewer pAVs are present in *itgb1a/b*-depleted embryo co-injected with 200 pg wild-type *itgb1* mRNA. No flow-carrying pAVs are apparent in rescue embryo, indicating a successful rescue. Flow-carrying AVs are again apparent in *itgb1a/b*-depleted embryo co-injected with 200 pg *itgb1* Δ10 mRNA, suggesting a failure to rescue.

A total of 46 LoF and 72 D-mis total damaging mutations were identified in genes included under axon guidance terms REAC:R-hsa-422475 (see above) or related axon guidance term GO:0007411 (p=2.16×10^-8^), including five DNVs (**Table S11**). Besides *RASA1* and *EPHB4*, several other high-pLI genes with essential roles in cerebrovascular development contained multiple rare, damaging *de novo* or transmitted variants, including *ACVRL1*, *NOTCH1*, *PTPN11*, and *ITGB1* (**Table S12**). These genes also contributed to enrichment signals identified in both the heart development term GO:0007507 and/or the Ras GTPase activator activity term GO:0005096 (**Fig. 2a**) and mapped onto a robust STRING ^70^ protein-protein interactome (PPI enrichment p-value < 1.0×10^-16^) with other Ras signaling genes harboring rare, damaging variants (**Table S13** and **Fig. 2b**). We therefore examined the probands harboring these variants in greater detail.

KVOGM73-1 harbored a novel *de novo* p.Gly616Cys variant in *NOTCH1* (**Table S12 and Fig. S3**), encoding a cell surface receptor for Jagged-1 (*JAG1*) and other Notch family ligands ^71^. *NOTCH1* mutations are associated with congenital heart defects and other vascular anomalies in autosomal dominant Aortic valve disease type 1 (OMIM# 109730) ^72^ and autosomal dominant Adams-Oliver syndrome type 5 (OMIM# 616028) ^73^. Although KVOGM73-1 exhibited no structural cardiac pathology, he had moyamoya disease (MMD), in common with *JAG1*-mutant patients with autosomal dominant Alagille syndrome type 1 ^74^ (OMIM# 118450). Notch proteins are characterized by N-terminal epidermal growth factor (EGF)-like repeats followed by LNR domains which form a complex with ligands to prevent signaling. p.Gly616Cys impacts a conserved residue in NOTCH1 required for proper folding of the EGF-like domain (**Fig. S3**) and is expected to activate Ras/ERK/MAPK signaling ^75^. Unrelated proband KVOGM83-1 contained another rare, damaging unphased mutation in *NOTCH1* (p. Asp1064Asn) at the surface exposed residue of the 28^th^ EGF repeat, which is part of the Abruptex region in fly Notch and potentially implicated in cis-inhibition of Notch (**Fig. S3**) ^76^. To our knowledge, this is the first report of co-existing VOGM and MMD, as well as the first report of a *de novo NOTCH1* mutation in a VOGM patient.

Proband KVOGM23-1 harbored a *de novo* p.Tyr63Cys variant in *PTPN11*, encoding the non-receptor tyrosine phosphatase SHP2 (**Table S12** and **Fig. S3**). *PTPN11* mutations, including the identical p.Tyr63Cys variant, have been reported in autosomal dominant Noonan syndrome type 1 (OMIM# 163950), which features a broad spectrum of congenital heart defects and other systemic vascular lesions ^77, 78^. KVOGM23-1 exhibited no Noonan syndrome-like features. Interestingly, p.Tyr63Cys impacts a conserved residue of SHP2’s N-terminal SH2 domain (**Fig. S3**). Similar to other N-SH2 domain ‘blocking loop’ variants, p.Tyr63Cys disrupts SHP2 autoinhibition and causes constitutive Ras signaling activation ^79, 80^. The paralogous p.Asp61Gly variant in SHP1 potentiates Ras/ERK/MAPK signaling by interfering with the phosphorylation of GAB1, an adaptor protein recruiting p120 RasGAP to receptor tyrosine kinases ^79, 81, 82^. To our knowledge, this is the first report of a *de novo PTPN11* variant in a VOGM patient.

Proband KVOGM-91 harbored a rare, damaging variant in *ACVRL1* (p.Cys344Tyr) (**Fig. 2c and Fig. S3**). Variants in *ACVRL1* encoding ALK1, a type I cell-surface TGF-beta superfamily receptor serine/threonine kinase ^83^, have been implicated in autosomal dominant hemorrhagic telangiectasia (HHT) type 2 (Rendu-Osler-Weber syndrome 2; OMIM# 600376). Whereas the proband had no features of HHT2, further inquiry revealed that his uncle died at two weeks of age from VOGM-related high-output heart failure. Moreover, the proband’s father, grandfather, and multiple paternal cousins had stigmata of HHT2, including epistaxis, visceral AVMs (lung and liver), and cutaneous vascular lesions. ALK1 p.Cys344Tyr segregated with VOGM and HHT2 vascular phenotypes in all affected family members available for sequencing. p.Cys344Tyr impacts a highly conserved residue in the kinase domain (**Fig. 2c**), has been identified in another HHT2 family (though without a history of VOGM) ^84^, and has been shown to decrease ALK1 cell surface expression ^85^.

In further support of the pathogenicity of *ACVLR1* mutation in VOGM, we found that *Acvrl1a/b* depletion in the *Tg(kdr:gfp)^zn^*^1^ reporter fish line (see **Methods**) resulted in VOGM-like massive dilation of the venous primordial hindbrain channel and posterior connecting segment, similar to that seen in *Ephb4a/b* deficient fish **(Fig. 2e)**; we also observed erratic connections between the venous posterior hindbrain channel and arterial basilar artery. These phenotypes were rescued by co-injection of wild-type human *Acvrl1* but not the VOGM-mutant human *Acvrl1* Cys344Tyr (**Fig. 2f and Fig. S7**).

Unrelated proband KVOGM-100 contained another rare, damaging variant in *ACVRL1* (p.Arg484Gln) (**Fig. 2c** and **Table S12**). KVOGM-100 had features of HHT2, including recurrent nosebleeds and pulmonary arterial hypertension (**Table S12**). The proband’s father carried the *ACVRL1* variant and had telangiectasias on his tongue, lips, and lower extremities. Variant p.Arg484Gln impacts a conserved surface-exposed residue located in helix αI of the kinase C-lobe, and encodes a catalytically inactive ALK1 mutant ^86^. Mutations in this region are associated with HHT2, an increased incidence of pulmonary arterial hypertension ^87–89^, and with other ALK family-associated diseases such as brachydactyly type A2 (ALK6) ^90, 91^ and Loeys-Dietz syndrome (ALK5) ^92, 93^. ACVRL1 p.Arg484Gln has also been identified in HHT2 ^94^ and in childhood-onset pulmonary arterial hypertension ^95^. p.Arg484Gln. ALK1 interacts with p120 RasGAP via the Dok-1 adaptor protein ^96^, implicating ALK1-DOK1-p120 RasGAP in crosstalk between the TGF-beta and Ras/ERK/MAPK signaling pathways.

KVOGM89-1 and KVOGM105-1 exomes respectively contained the rare, damaging, variants p.Ser785fs (unphased) and c.2331-2 A>G (transmitted) in *ITGB1* (Integrin subunit beta-1), which is essential for endothelial cell adhesion, migration, and survival during angiogenesis ^97^ (**Fig. S3 and Table S12)**. Interestingly, both patients had intracranial hemorrhage, progressive macrocephaly, and shunt-dependent hydrocephalus. KVOGM105-1 and his sister (unavailable for sequencing) presented with impressive nevus simplex lesions, a type of capillary malformation. p.Ser785fs and c.2331-2 A>G mutations are predicted to lead to nonsense-mediated RNA decay.

In support of the pathogenicity of these *ITGB1* variants in VOGM, *Itgb1a/b* depletion in the *Tg(kdr:gfp)^zn^*^1^ reporter fish line resulted in VOGM-like supernumerary connections between venous and arterial vessels similar to *Ephb4a/b* and *Acvrl1a/b* mutant fish (**Fig. 2g and Fig. S7**). Cerebral microangiography showed numerous abnormal high-flow connections between the venous primordial hindbrain channel and the arterial basilar artery (**Fig. 2g**; see above). Defects in *Itgb1a/b* were rescued by co-injection of wild-type human *Itgb1* but not by the VOGM-associated mutant human *Itgb1* Δ10.

Together, these results suggest that rare, damaging *de novo* or transmitted variants in other mutation-intolerant genes implicated in other vascular Mendelian syndromes and Ras signaling are pathogenic contributors to VOGM.

### VOGM risk genes converge in growth factor-regulated Ras/ERK/MAPK signaling networks in developing cerebral endothelial cells

To gain insight into the developmental periods, cell types, and molecular pathways involved in VOGM biology, we performed a spatiotemporal consensus Weighted Gene Co-expression Network Analysis (WGCNA) ^98^ leveraging a large bulk RNAseq data set encompassing multiple human brain regions across development and into early childhood ^99^. We constructed 88 modules characterized by genes that share highly similar expression patterns during brain development across different cortical regions and therefore likely to be involved in similar functions ^98^. Each module was assessed for relative enrichment of high-confidence and probable (p) VOGM genes, brain AVM genes, AVM+VOGM genes together (since each is a high-flow arterio-venous communication disorder), cerebral cavernous malformation (CCM) genes, moyamoya disease (MMD) genes, and human height genes (as a negative control) using multivariate logistic regression (see **Table S14** and **Methods** for description of gene lists).

VOGM genes converged in several modules in the fetal human cortex. The most significant of these was the post-conceptional week (PCW) 37 “Midnight Blue” module (P = 2.13 × 10^-6^; **Fig. 3**). The “Black” module was also enriched with VOGM genes (**Fig. 3a**). The “Black” module exhibited peak gene expression early in development at PCW 9-17 (**Fig. 3b**). In contrast, the only module to show enrichment for VOGM genes alone, “Light Cyan”, is expressed later in neurodevelopment, at 10 – 12 months postnatally. (**Fig. 3b**). Notably, genes expressed in the “Midnight Blue” module are enriched for those involved in Focal Adhesion-PI3K-Akt-mTOR Signaling Pathway (WP:3932; P = 6.73 × 10^−10^, 6.7-fold enrichment), including Positive Regulation of Vascular Development (GO: 1904018; enrichment 11.4-fold; P = 7.95 × 10^−8^) and Regulation of Cell Migration (GO: 0030334; enrichment 4.4-fold; P = 5.94 × 10^−6^; **Fig. 3c**). Human height genes were unenriched in any of the defined modules.

**Figure 3.**
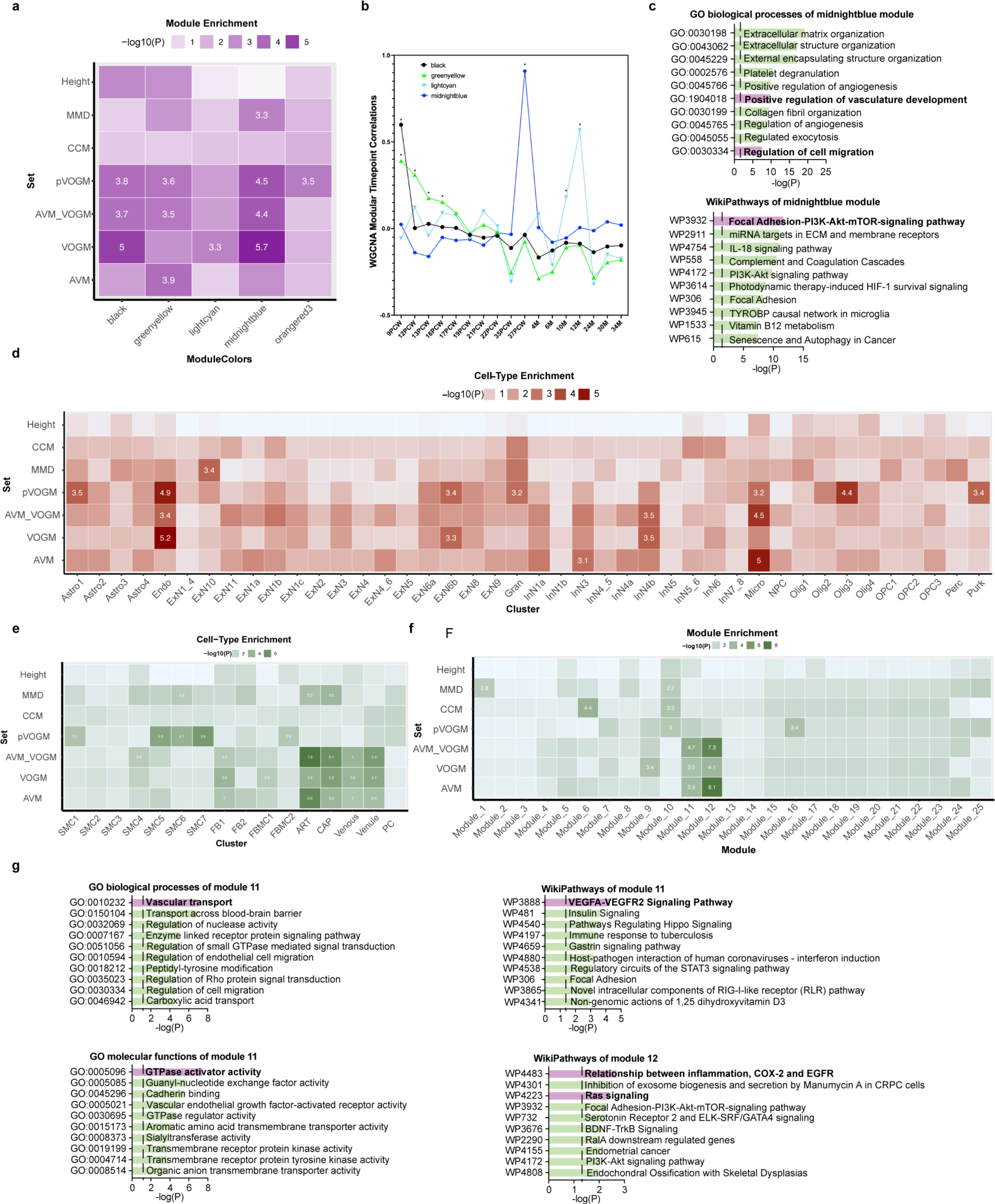
VOGM genes converge in a VEGFR-Ras signalling network in fetal cerebral endothelial cells. (a) Enrichment of VOGM genes in gene modules of the fetal human cortex compared to other disease genes. Numbers displayed exceed the Bonferroni-corrected statistical significance threshold and are −log_10_(p-value). Height: human height gene set; MMD: moyamoya disease gene set; CCM: cavernous malformation gene set; AVM_VOGM: arteriovenous malformation and vein of Galen aneurysmal malformation disease gene set; VOGM: high-confidence vein of Galen aneurysmal malformation disease gene set; pVOGM: probable VOGM gene set; AVM: arteriovenous malformation disease gene set (see Methods for gene set determination details). (b) Temporal dynamics of modules enriched with VOGM genes. The peak expression of “Midnight Blue” module is at post-conception week (PCW) 37. Both the “Black” and “Green-yellow” modules exhibited peak gene expression early in development at PCW 9-17. “Light Cyan” module is expressed much later in neurodevelopment at postnatal age 10 – 12 months. (c) Gene Ontology (GO) biological processes and WikiPathways of midnight blue module converge on Focal Adhesion-PI3K-Akt-mTOR Signaling Pathway, Positive Regulation of Vascular Development, and Regulation of Cell Migration. The significance threshold, adjusted for multiple comparisons, is denoted by the vertical dashed line. Top enriched terms have bolded text and purple bars. (d) Cell-type enrichment of VOGM and other disease genes in the fetal human cortex defined by scRNAseq. Numbers displayed exceed the Bonferroni-corrected statistical significance threshold and are - log_10_(p-value). Different cell types are noted on the x-axis, see text for details. (e) Cell-type enrichment of VOGM genes in the developing human cerebrovasculature defined by scRNAseq. Numbers displayed exceed the Bonferroni-corrected statistical significance threshold and are −log_10_(p-value). Different cell types are noted on the x-axis (see text for details). (f) Enrichment of disease genes in fetal human cortex modules. Numbers displayed exceed the Bonferroni-corrected statistical significance threshold and are −log_10_(p-value). (g) GO molecular function and WikiPathways analyses of enriched module 11 and 12. The significance threshold, adjusted for multiple comparisons, is denoted by the vertical dashed line. Enriched terms of interest have bolded text and purple bars.

We next studied VOGM gene expression using two different single cell (sc)-RNAseq transcriptomic atlases. The first set comprised of 154,938 total cells across 42 cell types from 20 brain regions spanning different developmental stages ^100^. VOGM genes (P = 6.43 × 10^-6^) and pVOGM genes (P=1.25 x 10^-5^) showed a specific enrichment in the endothelial cell (EC) subcluster. VOGM+AVM genes were also enriched in ECs, but the contributors to this signal were VOGM-specific genes as AVM genes alone did not yield significant enrichment. In contrast, CCM and MMD genes were not enriched in ECs. VOGM also showed enrichment in other cell types to lesser degrees of enrichment than in EC subtypes. Examples include the excitatory embryonic neuron (cluster “ExN6b”, P = 4.75 × 10^-4^) and the inhibitory neurons (cluster “InN4b”, P = 3.18× 10^-4^) (**Fig. 3d**). Control human height genes were unenriched in any of the 42 cell subtypes.

The second sc-RNAseq transcriptomic atlas included 181,388 total cells from normal and diseased human cerebrovasculature, including both endothelial and perivascular cell subtypes ^46^. Corroborating results from the STAB dataset, VOGM genes showed a specific enrichment in ECs (P = 6.43 x 10^-6^). VOGM, AVM, and AVM+VOGM genes were enriched in arterial and venous EC subtypes, but not in perivascular subtypes such as smooth muscle cells, fibroblasts, fibromyocytes, pericytes, or other perivascular cell types (**Fig. 3E**). In contrast, CCM and MMD genes were unenriched in arterial and venous EC subtypes. Human height genes were not enriched in any of the 18 cell subtypes.

We again constructed modules from the cerebrovascular dataset comprising co-expressed genes. These were then assessed for the enrichment of VOGM and other disease genes (see **Methods**). Of the 25 defined modules, VOGM genes were most highly enriched in Modules 11 (P = 3.23 × 10^-4^) and 12 (P = 8.22 × 10^-5^), overlapping with AVM genes (**Fig. 3f**). In contrast, CCM and MMD genes were most enriched in Module 6 defined by mitochondrial processes. Human height genes were not significantly enriched in any of the Modules (**Fig. 3f**). Module 11 genes were most highly enriched for pathways related to VEGFA-VEGFR2 signaling (WP3888; 1.97-fold enrichment; P = 4.51 x 10^-5^), vascular transport (GO:0010232; 5.0-fold enrichment; P = 7.04 x 10^-8^), and GTPase activator activity (GO:0005096; 2.7-fold enrichment; P = 3.89 x 10^-8^) (**Fig. 3g and Fig. S8**). Module 12 was notably enriched for pathways related to Ras/ERK/MAPK (WP4223; 2.3-fold enrichment; P = 3.89 x 10^-3^) and inflammation-regulated EGFR signaling (WP4483; 6.5-fold enrichment; P = 2.05 x 10^-3^) (**Fig. 3g and Fig. S8**). Taken together, these results implicate growth factor-regulated Ras/ERK/MAPK signaling in fetal cerebral endothelial cells as an important spatio-temporal locus impacted by VOGM-associated gene mutations.

### “Two-hit” VOGM-specific *EPHB4* kinase domain missense mutation activates Ras/ERK/MAPK signaling in fetal endothelial cells and disrupts developmental angiogenesis

Mice constitutively deficient in EPHB4 or its ligand, Ephrin B2, exhibit blood vascular abnormalities and expire at E10.5 of gestation because vascular plexuses that arise through vasculogenesis are not remodeled by angiogenesis into hierarchical arterial-capillary-venous networks ^3,101, 102^. However, the possible requirement of EPHB4 kinase activity for normal vascular development is unknown. Based on our human genetic (**Fig. 1**) and integrative genomic results (**Fig. 3**), we hypothesized that genetic inactivation of a kinase-dependent, EPHB4-regulated RASA1 signaling mechanism in endothelial cells causes constitutive Ras-MAPK activation and disruption of VEGF-associated developmental angiogenesis *in vivo*. To test this hypothesis and gain insight into the mechanism of the identified VOGM-associated EPHB4 kinase domain missense mutations, we generated *EphB4* mutant mice carrying a knock-in allele orthologous to the *EPHB4* p.Phe867Leu variant identified in patient KVOGM18-1 (**Table 2** and **Methods**).

Heterozygous *Ephb4^+/F867L^* mice were viable, fertile, and exhibited no obvious vascular developmental or other defects. However, homozygous *Ephb4^F867L/F867L^* pups were not identified among the progeny of heterozygous *Ephb4^+/F867L^* intercrosses, suggesting that the EPHB4 p.Phe867Leu variant is lethal in homozygous form. Examination of embryos at E10.5 confirmed striking vascular defects in *Ephb4^F867L/F867L^* embryos similar to those reported in constitutive EPHB4-and Ephrin B2-deficient E10.5 embryos (**Fig. 4a**). Blood flow was absent in E10.5 *Ephb4^F867L/F867L^* embryos and the pericardial sac was grossly distended. The yolk sacs of E10.5 *Ephb4^+/+^* and *Ephb4^+/F867L^* embryos showed normal hierarchical vascular networks supplying blood to the developing embryo (**Fig. 4a and b**). In contrast, the yolk sacs of E10.5 *Ephb4^F867L/F867L^* embryos exhibited honeycomb-like vascular patterning typical of primitive vascular networks (**Fig. 4a and b**). At E10.5,

**Figure 4.**
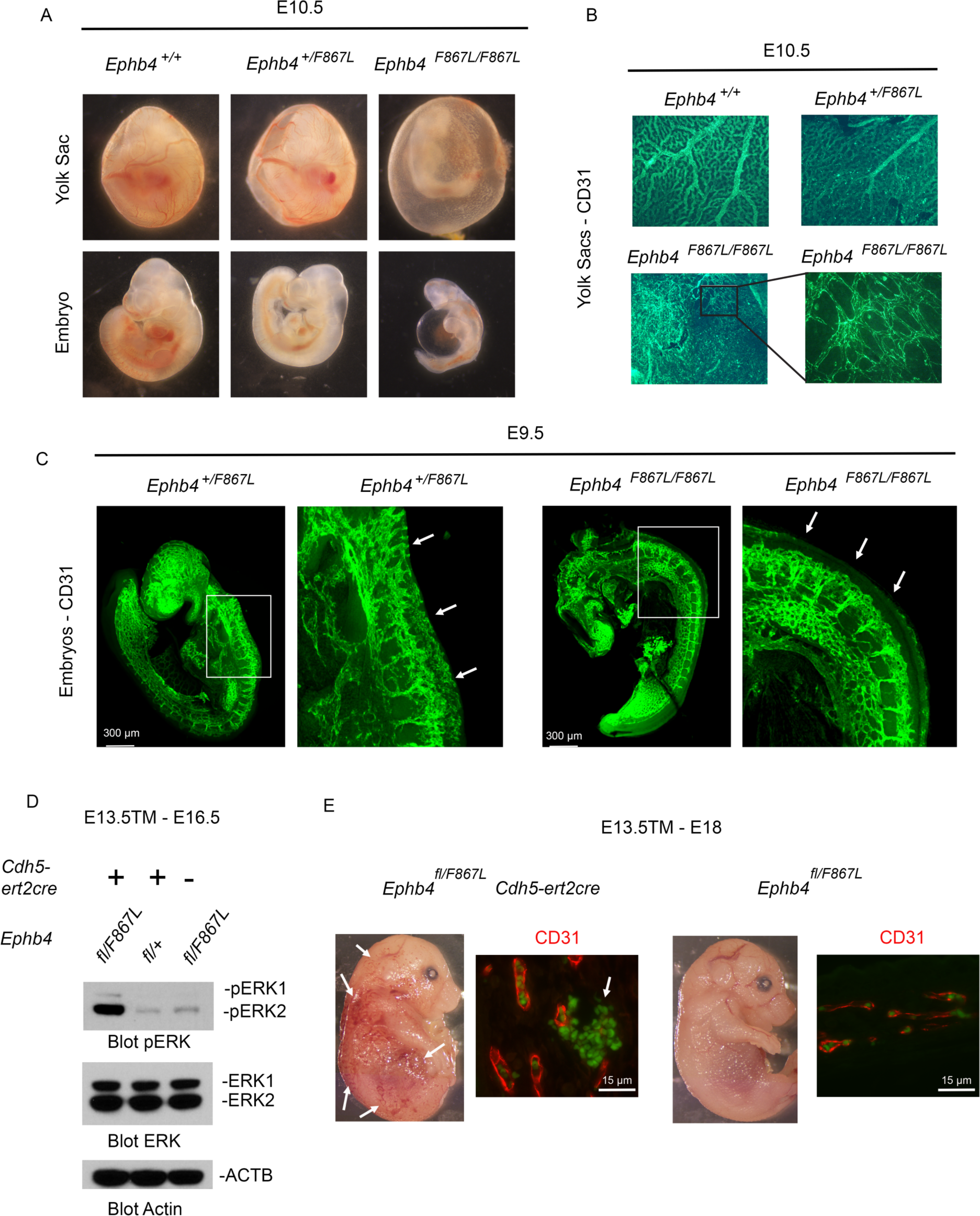
VOGM-specific *EPHB4* mutation activates endothelial Ras-MAPK signaling and disrupts developmental angiogenesis. (a) Vascular defects in homozygous and heterozygous EPHB4 mutant embryos. Shown are images of littermate embryos of the indicated genotypes at E10.5. Note abnormal vascularization of the yolk sac, reduced size and distended pericardial sac of homozygous EPHB4 F867L mouse embryos. (b) Honeycomb-like vascular patterning in homozygous F867L EPHB4 mutants. E10.5 yolk sacs were stained with anti-CD31 antibodies to identify the vasculature. For the EPHB4 F867L yolk sac, the image right is a higher magnification of the boxed area in the left image. Note hierarchical vascular networks in WT and EPHB4 F867L heterozygous yolk sacs and primitive vascular plexuses in EPHB4 F867L homozygous yolk sacs. (c) Characterization of vascular defects E9.5 *Ephb4^F^*^867^*^L/F867L^* embryos. CD31 antibody staining of littermate E9.5 embryos of the indicated genotypes. For each embryo, the right image is a higher magnification of the boxed area in the left image. Arrows show trunk angiogenesis toward the midline in EPHB4 F867L heterozygous but not EPHB4 homozygous embryos. (d and e) Embryos of the indicated genotypes were administered tamoxifen at E13.5 and harvested at the indicated time points. (d) MAPK activation assessment of E16.5 embryos by Western blotting. Activation of MAPK in liver lysates was determined by Western blotting using anti-pERK antibodies. Equal protein loading was determined by antibody probing for ERK and actin abundance. (e) Extensive cutaneous vascular hemorrhage in *Ephb4^fl/F867L^ Cdh5ert2cre* embryos. Gross images of E18 embryos (left) and images of tissue sections of skin from the same embryos stained with anti-CD31 antibody (right). Note hemorrhagic appearance and extravasation of green auto-fluorescent erythrocytes in *Ephb4^fl/F867L^ Cdh5ert2cre* embryos (arrows).

*Ephb4^F867L/F867L^* embryos were reduced in size compared to *Ephb4^+/+^* and *Ephb4^+/F867L^* embryos, and histological examination revealed evidence of extensive apoptotic cell death **(Fig. 4a)**. Therefore, we examined E9.5 *Ephb4^F867L/F867L^* embryos to characterize vascular defects in the embryo. E9.5 *Ephb4^F867L/F867L^* embryos exhibited impaired developmental angiogenesis among intersomitic vessels that failed to extend branches toward the midline as observed in *Ephb4^+/F867L^* embryos (**Fig. 4c**). The same phenotype was previously noted in constitutive EphB4-and Ephrin B2-deficient embryos at E9.5 ^102^.

Whereas EPHB4 acts in transformed cells as a growth factor receptor that can activate Ras signaling, in endothelial cells, EPHB4 acts as an inhibitor of Ras signaling through functional interaction with RASA1 ^66, 103, 104^. Loss of kinase activity associated with VOGM-associated EPHB4 mutants could therefore dysregulate Ras activity in embryonic endothelial cells. To examine if endothelial-specific expression of EphB4 Phe867Leu EphB4 during this period of embryonic dysregulated Ras/ERK/MAPK signaling, we generated *Ephb4^fl/F867L^* embryos that carried the *Cdh5-ert2cre* endothelial-specific Cre driver (see **Methods**). At E13.5 of development, tamoxifen was administered to embryos to disrupt the *Ephb4* floxed allele resulting in the expression of EphB4 Phe867Leu exclusively in endothelial cells. At E16.5, embryos were harvested and Ras/ERK/MAPK activation in liver lysates was assessed by Western blotting using a phospho-specific antibody directed against ERK (**Fig. 4d)**. Compared to tamoxifen-treated *Ephb4^fl/+^ Cdh5-ert2^cre^* embryos and *Ephb4^fl/F867L^* embryos without a *cre* transgene, *Ephb4^fl/F867L^ Cdh5-ert2^cre^* embryos showed striking Ras/ERK/MAPK activation in the vasculature. By E18, *Ephb4^fl/F867L^ Cdh5-ert2^cre^* embryos showed evidence of extensive cutaneous vascular hemorrhage confirmed by CD31 immunostaining of embryo sections (**Fig. 4e**). These findings suggest VOGM-associated missense variants in *EPHB4* disrupt a kinase-dependent EPHB4-RASA1 signaling axis that normally limits activation of Ras/ERK/MAPK signaling in endothelial cells, which is important for angiogenic events required for normal hierarchical development of arterial-capillary-venous networks.

## DISCUSSION

The genetic understanding of VOGM has been hindered by its rarity and sporadic nature. Here, we presented the largest exome-sequenced cohort of VOGM to date and integrated these WES results with human brain and vasculature transcriptomes to define the cell types, developmental time points, and functional networks impacted by VOGM genes. We also validated new candidate genes in novel zebrafish and mouse models of disease (see **Graphical Abstract**). Collectively, these data shed insight into the cellular and molecular pathogenesis of VOGM, help illuminate the genetic regulation of human arterio-venous development, and have implications for clinical risk assessment of patients and their families and the development of targeted therapeutics.

Our analysis uncovered enrichment of rare, heterozygous, damaging *de novo* and inherited germline variants in VOGM probands in mutation-intolerant (high-pLI) genes, multiple of which are Ras/ERK/MAPK signaling genes (**Table S13 and S15**) that are highly expressed in the embryonic vasculature ^105^, are known regulators of multiple aspects of neurovascular development ^106, 107^, and map onto a robust STRING ^70^ protein-protein interactome (**Fig. 2**). Among these, p120 RasGAP (*RASA1*), harbored a genome-wide significant burden of *de novo* loss-of-function variants. Transmitted D-mis variants causing loss of receptor tyrosine kinase activity were enriched in EphB4 (*EPHB4*), which forms a complex with p120 RasGAP to negatively regulate Ras/MAPK/ERK signaling. The detected variants in each are shown *in vitro* or *in vivo*, or predicted *in silico*, to constitutively activate Ras/MAPK/ERK signaling.

Other probands had *de novo* or inherited pathogenic germline variants in other Ras-associated genes, such as a gain-of-function variant in the tyrosine phosphatase SHP2 (*PTNPN11*), which binds to and dephosphorylates Ras to increase its association with Raf and activate Ras/ERK/MAPK signaling ^108, 109^. Loss-of-function variants in the serine/threonine receptor kinase ALK1 (*ACVRL1*), which facilitates crosstalk between the TGF-beta and Ras/ERK/MAPK signaling pathways by associating with p120 RasGAP via the Dok-1 adaptor protein ^96^, were detected in other probands, including an ultra-rare multi-generational VOGM family. These data suggest that genetically-encoded disinhibition of Ras/ERK/MAPK signaling is an important driver of VOGM pathogenesis, and expand the spectrum of the cerebral vascular anomalies associated with the Ras-opathies ^25, 27–29, 110–113^ (see **Graphical Abstract**).

Germline mutations in VOGM risk genes *RASA1*, *EPHB4*, *ACVRL1*, *PTNPN11*, and *NOTCH1* have been reported in other Mendelian diseases featuring vascular phenotypes ^25, 27, 109, 110, 114–116^*. RASA1* and *EPHB4* encode interacting proteins mutated in CM-AVM type 1 and 2, respectively. Our VOGM probands with *ACVRL1* (ALK1) variants, which have been linked with HHT1, are either part of a multi-generational VOGM family with HHT1 or have HHT1 features. Consistent with this, a VOGM patient with a mutation in HHT2 gene *ENG*, encoding the ALK1 binding partner Endoglin, had been previously identified ^28^. Interestingly, the Ras-activating *PTPN11* (SHP2) p.Tyr63Cys variant identified in KVOGM23-1 has been previously identified in Noonan syndrome (OMIM# 163950), which features a broad spectrum of congenital heart defects and other systemic vascular lesions ^77, 78^. These results suggest VOGMs may represent phenotypic expansions of CM-AVM, HHT, or other Mendelian vascular syndromes. Nonetheless, it is interesting that none of our *RASA1* or *EPHB4* patients or their family members carried a diagnosis of CM-AVM at the time of DNA sequencing.

Inherited VOGM mutations show incomplete penetrance and variable expressivity, with mutation carriers often exhibiting cutaneous vascular lesions. Cutaneous vascular lesions are a common hallmark of developmental vascular disorders such as *RASA1*-and *EPHB4*-mutated CM-AVM ^25, 27^, *ENG1*-and *ACVRL1*-mutated HHT ^29^, and *RASA1*-mutated Parkes Weber syndrome, among others ^30, 117, 118^. As many VOGM families with full clinical data had capillary malformations or other uncommon cutaneous vascular lesions, and as mutations identified in probands were found in all family members with these cutaneous lesions, VOGMs and the cutaneous lesions are likely linked to the same mutations. These observations highlight the pleiotropy of these gene mutations and the value of WES as a diagnostic adjunct.

Variable expressivity of VOGM and associated features could arise from environmental modifiers ^119^ in concert with the identified rare mutations and/or specific genetic modifiers ^43^. However, our findings are most consistent with a two-hit mechanism in which phenotypic expression relies on an inherited germline mutation and a second, post-zygotic (i.e., somatic) mutation in the other wild-type allele ^120, 121^. This mechanism has been shown for other hereditary multifocal vascular malformations, such as CM-AVM1 ^27, 122, 123^, glomuvenous malformations (OMIM: 138000), cutaneomucosal venous malformation (OMIM: 600195), and cerebral cavernous malformations (OMIM: 116860) ^121^. In this context, phenotypic expression depends on the cell types in which somatic mutations co-occur and could explain the low penetrance of VOGM arising from transmitted mutations. Our data in mice supports this genetic mechanism, as heterozygous *Ephb4^F867L^* mice expressing a VOGM-specific *EPHB4* kinase-domain missense variant exhibited severe vascular abnormalities, but only when carrying a second variant allele. Exome sequencing of ultra-rare lesional human VOGM tissue could test this two-hit hypothesis.

Integration of our WES findings with scRNAseq datasets of the developing human cerebrovasculature helped define developing endothelial cells as a critical spatio-temporal locus of VOGM pathophysiology. Pathway analysis suggested that, in developing endothelial cells, VOGM genes play critical roles in growth-factor-regulated, tyrosine receptor kinase-associated Ras/MAPK/ERK signaling, which is known to regulate multiple aspects of vascular morphogenesis including vasculogenesis, angiogenesis, and arterio-venous specification ^1^. Consistent with this, tamoxifen-induced *Ephb4^fl/F867L^ Cdh5ert2cre* mice expressing a VOGM-patient mutation that disrupts EphB4 kinase activity in fetal endothelial cells exhibited impaired remodeling of primitive vascular plexuses by VEGF-regulated sprouting angiogenesis, which is required for the development of hierarchical arterial-capillary-venous networks ^66^. VOGM genes are also enriched in pathways that regulate axon pathfinding, which is known to be important for vascular patterning ^1^. Further mechanistic details of how VOGM mutations impact vasculogenesis, including the intercellular communication between developing neural and endothelial cell types, will be important topics of future investigation.

Although rare, damaging mutations of large effect contribute ∼10.3% of VOGM cases, mutations in additional genes likely contribute to disease pathogenesis. Our Monte Carlo simulation based on observed damaging DNVs estimated ∼66 genes contribute to VOGM by a *de novo* mechanism. WES of 250 or of 1000 trios are expected to yield respective saturation rates of 20.1% and 70.5%, respectively (**Fig. S9**). The consequence of cis-regulatory elements and more complex structural variants remains unknown. The still unclear impact of somatic mutations requires additional deep sequencing in matched normal and affected-tissue pairs to enable their characterization. Genomic repeats, transposable elements, and epigenomic changes should also be investigated using the more comprehensive long-read sequencing technologies.

Our findings have several clinical implications. The enrichment of damaging variants in the VOGM cohort suggests that mutation carrier offspring may be at increased risk for VOGMs as well as for capillary malformations and potentially other AVMs. However, not all variant carriers develop capillary malformations, making the presence of capillary malformations an unreliable clinical marker for transmission risk in affected families. These observations highlight the importance of family history and the potential use of exome sequencing-based screening for risk assessment among family members. Family members of VOGM patients with positive exome sequencing results could possibly benefit from intracranial imaging with MRI/MRA, especially in the setting of suspicious mucocutaneous lesions.

MAPK inhibitors rescue *RASA1* and *EPHB4*-mutant embryonic vascular phenotypes in mice ^66, 124, 125^. We have shown that a VOGM-associated *EPHB4* missense mutation causes constitutive Ras/MAPK/ERK activation in developing endothelial cells and impaired hierarchical development of angiogenesis-regulated arterial-capillary-venous networks. Therapies targeting the MAPK pathway may therefore represent a viable therapeutic approach for VOGM and, perhaps other CM-AVM and HHT spectrum lesions ^66, 124, 125^. Trametinib, an orally available inhibitor of MEK kinase activity, could be a potential treatment for VOGM and other vascular anomalies involving the pathological activation of Ras/MAPK/ERK signaling ^126, 127^. Indeed, a prospective phase II trial, TRAMAV (Evaluating the safety and efficacy of Trametinib in Arterio-Venous Malformations that are refractory to standard care) is currently ongoing (https://www.clinicaltrialsreg-ister.eu; Unique identifier: 2019-003573-26). However, in contrast to systemic vascular lesions, the narrow window of gestational weeks 6–11 during which the primitive choroidal arteries and the MPV are usually present, and therefore during which VOGMs are speculated to develop^10^, may pose a challenge to improved early therapeutic strategies for VOGM. Thus, attempted diagnosis with the intention to treat would have to occur before the safe gestational age threshold for amniocentesis ^128^. These data highlight the need for continued genetic research on VOGM, with a focus on the mechanistic implications of recently discovered VOGM-associated variants in mammalian model systems.

## ACKNOWLEDGEMENTS

This work is supported by the Yale-National Institutes of Health (NIH) Center for Mendelian Genomics (5U54HG006504); R01 NS111029-01A1, R01 NS109358, K12 228168, and the Rudi Schulte Research Institute (K.T.K.); R01 117609 (K.K. and T.J.B.); NIH Medical Scientist Training Program (NIH/National Institute of General Medical Sciences Grant T32GM007205); NIH Clinical and Translational Science Award from the National Center for Advancing Translational Science (TL1 TR001864); the K99/R00 Pathway to Independence Award (K99HL143036 and R00HL143036-02), the Vernon W. Lippard Research Fellowship, the Hydrocephalus Foundation Innovator Award (K.T.K. and S.C.J.), the March of Dimes, the Howard Hughes Medical Institute, R01 HL146352 and 2R01 HL120888 (both P.D.K.), and WashU Clinical & Translational Research Funding Program award (CTSA1405), and WashU Children’s Discovery Institute Faculty Scholar award (CDI-FR-2021-926).

## AUTHOR CONTRIBUTIONS

**Study design and conceptualization:** K.T.K., S.C.J., S.Z, K.Y.M., P.D.K.

**Cohort ascertainment, recruitment, and phenotypic characterization:** K.Y.M, H.S., J.O., J.S., A.J.K., T.D.J., A.B.W.G., P.P., B.J.I., B.A-K., A.T.H., J.M.J., E.J., P.B.S., S-S.L., W.E.B., B.S.C., P.C., C.J.S., A.B.P., G.R., S.S., A.B., A.P.S, T.B., J.Z., K.P.F., J.E.C., M.R.P., E.R.S., M.G., R.P.L., S.C.J., K.T.K.

**Exome sequencing production and validation:** S.M., R.P.L.

**Exome sequencing analysis:** S.Z., X.Z., W.D., D.D., C.G.F., B.C.R., P-Y. F., Y-C. W.

**Integrative genomics analysis:** G.A., S.Z., S.C.J., K.T.K.

**Phenomics analysis:** K.Y.M., M.R., J.S., S.Z. J.O., H.S., A.J.K., T.D.J., S.K., S.M.K.

**Statistical analysis:** S.Z., K.Y.M, G.A., S.C.J., K.T.K. B.L., H.Z., E.Z.E-O.

**Sanger sequencing validation:** C.N-W.

**Neuroimaging characterization:** K.T.K., D.B.O., A.M.-D.-L.

**Structural analysis:** J.E.C., T.J.B.

**Biophysical simulation:** S.Z.

**Functional experimentation and analysis:** M.A.vdE., A.P., P.Q.D., D.C., S.N., T.N., S.B., H.T.L., S.Z., K.T.K.

**Resources:** K.T.K., S.C.J., P.D.K.

**Writing and review of manuscript:** K.T.K., S.C.J., P.D.K., S.Z., K.Y.M., M.A.vdE., G.A., A.P., T.J.B., S.L.A., S.N., W.E.B., D.B.O., S.K., S.M.K.

**Project administration:** K.T.K., S.C.J., P.D.K., S.Z., K.Y.M.

**Funding acquisition and supervision:** K.T.K., S.C.J., P.D.K., T.J.B.

## METHODS

### Patient subjects

All procedures in this study comply with Yale University’s Human Investigation Committee (HIC) and are approved by Yale University’s Human Research Protection Program. Written informed consent was obtained from all adult participants. Written authorization was obtained from a parent or legal guardian for sample collection from all minors in this study. Inclusion criteria included male or female patients with clearly defined mural or choroidal VOGMs, radiographically confirmed by both a neurosurgeon and neuroradiologist from an angiogram or magnetic resonance angiogram. Family members of included patients were also studied when possible. Controls consisted of 1,798 unaffected siblings of autism cases and unaffected parents from the Simons Foundation Autism Research Initiative Simplex Collection (SSC) ^49, 50^. Only the unaffected siblings and parents, as designated by SSC, were included in the analysis and served as controls for this study. Permission to access the genomic data in the SCC or the National Institute of Mental Health Data Repository was obtained. The Simons Foundation Autism Research Initiative provided written informed consent for all participants.

### Exome sequencing and variant calling

Exome capture was performed on genomic DNA samples derived from saliva or blood using Roche SeqCap EZ MedExome Target Enrichment kit or IDT xGen target capture kit followed by 99 base paired-end sequencing on the Illumina sequencing platform. Sequence reads were aligned to the human reference genome GRCh37/hg19 using BWA-MEM ^129^ and further processed to call variants following the GATK Best Practices workflow ^51^ and Freebayes ^53^. Variants were annotated with ANNOVAR, gnomAD (v.2.1.1), and Bravo databases ^54^, Karczewski, et al. ^130, 131^. We used MetaSVM and MPC algorithms to predict the deleteriousness of missense variants (D-Mis, defined as MetaSVM-deleterious or MPC-score ≥ 2) ^56, 132^. Inferred loss-of-function (LoF) variants consist of stop-gain, stop-loss, frameshift insertions/deletions, canonical splice sites, and start-loss. LoF and D-Mis mutations were considered ‘damaging.’ *De novo* variants (DNVs) were called using TrioDeNovo ^55^. Candidate DNVs were further filtered based on the following criteria: (1) exonic or splice-site variants; (2) a minimum read depth (DP) of 10 in the proband and both parents; (3) minimum proband alternative read depth of 5; (4) proband alternative allele ratio ≥ 28% if having < 10 alternative reads or ≥ 20% if having ≥ 10 alternative reads; (5) alternative allele ratio in both parents ≤ 3.5%; (6) global MAF ≤ 4 x 10^-4^ in the Exome Aggregation Consortium database. For recessive variant analysis, we filtered for rare (MAF ≤ 1 × 10^−3^ in Bravo and in-cohort MAF ≤ 1 × 10^−2^) homozygous and compound heterozygous variants that exhibited high-quality sequence reads (pass GATK variant quality score recalibration, ≥ 8 total reads for proband, genotype quality (GQ) score ≥ 20). Only LoF, D-Mis, and non-frameshift indels were considered potentially deleterious to the disease. Only homozygous variants were analyzed for probands with unavailable parental WES data. See **Table S16** for damaging recessive variants identified. For rare transmitted dominant variants, only LoF and D-Mis were included. They were filtered using the following criteria: (1) pass GATK variant quality score recalibration, (2) MAF ≤ 5 × 10^−5^ in Bravo and in-cohort MAF ≤ 5× 10^-3^; (3) a minimum DP of 8 in the proband, and (4) GQ score ≥ 20. After filtering each type of mutation as described above, false-positive calls were removed by *in silico* visualization. Candidate mutations were confirmed by PCR amplification followed by Sanger sequencing (primer sequences available on request).

### Kinship analysis

The relationship between proband and parents was estimated using the pairwise identity-by-descent (IBD) calculation in PLINK ^133^. IBD sharing between the proband and parents in all trios was between 45% and 55%. For pairs sharing ≥ 80% of rare variants, the sample with greater sequence coverage was retained in the analysis, and the other discarded.

### Principal component analysis

Subject ethnicity was determined by EIGENSTRAT ^134^ software to analyze tag SNPs in cases, controls, and HapMap subjects, as described ^39^.

### *De novo* expectation model

As case trios were captured by two different reagents (MedExome and IDT), we took the union of all bases covered by different capture reagents. We generated a Browser Extensible Data file representing a unified capture for all trios. We used bedtools (v.2.27.1) to extract sequences from the Browser Extensible Data file. We then applied a sequence context-based method to calculate the probability of observing a DNV for each base in the coding region, adjusting for sequencing depth in each gene as described previously ^135^. Briefly, for each base in the exome, the probability of observing every trinucleotide mutating to other trinucleotides was determined. ANNOVAR (v2015Mar22) was used to annotate the consequence of each possible substitution. RefSeq was used to annotate variants (based on the file ‘hg19_refGene.txt’ provided by ANNOVAR). For each gene, the coding consequence of each potential substitution was summed for each functional class (synonymous, missense, canonical splice site, frameshift insertions/deletions, stop-gain, stop-loss, and start-loss) to determine gene-specific mutation probabilities ^135^. The probability of a frameshift mutation was determined by multiplying the probability of a stop-gain mutation by 1.25, as described previously ^135^. The model does not account for in-frame insertions or deletions. To align with ANNOVAR annotations, analysis was limited to variants located in exonic or canonical splice site regions and were not annotated as ‘unknown’ by ANNOVAR. We identified potential coding mutations and generated gene-specific mutation probabilities for 19,347 unique genes following the inclusion criteria. Due to the difference between exome capture kits and DNA sequencing platforms, and to variable sequencing coverage between case and control cohorts, separate de novo probability tables were generated for cases and 1,798 autism control trios.

### *De novo* enrichment analysis

The burden of DNVs in the case and control trios was determined using the denovolyzeR package as previously described ^39, 136^. Briefly, the expected number of DNVs in the case and control cohorts across each functional class was calculated by taking the sum of each functional class-specific probability multiplied by the number of probands in the study 2× (diploid genomes). Then, the expected number of DNVs across functional classes was compared to the observed number in each study using a one-tailed Poisson test. Gene set enrichment analyses considered only mutations observed or expected in genes within the specified gene set (chromatin modifiers [GO:0016569] and LoF-intolerant). To examine whether any individual gene contains a greater number of protein-altering DNVs than expected, the expected number of protein-altering DNVs was calculated from the corresponding probability, adjusting for cohort size. A one-tailed Poisson test was then used to compare the observed DNVs for each gene versus expected. As separate tests were performed for protein-altering, protein-damaging, and LoF DNVs, the Bonferroni multiple-testing threshold is, therefore, equal to 8.6 × 10^−7^ (= 0.05 / (3 tests × 19,347 genes)).

### Estimation of the expected number of rare, transmitted variants

We implemented a one-tailed binomial test to quantify enrichment of rare damaging or LoF heterozygous mutations in each gene by comparing observed and expected counts estimated as previously ^39^. Because the number of rare damaging or LoF heterozygous mutations in a gene was inversely correlated with the constraint score (i.e., pLI score) obtained from the gnomAD database, we stratified genes into five subsets by pLI quartiles to control for the potential confounding effect: (1) the first quartile with pLI < 6.4 × 10^−8^; (2) the second quartile with pLI score 6.4 × 10^−8^ - 1.9 × 10^−3^; (3) the third quartile with pLI 1.9 x 10^-3^ - 0.48; (4) the fourth quartile with pLI 0.48 - 1.0; between third quantile and 1; and (5) those without a pLI score. For each set, the expected number of LoF heterozygous variants per gene was estimated by the following formula:

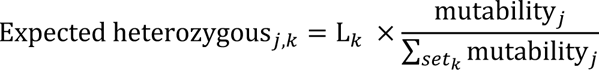

where ‘j’ denotes the ‘jth’ gene, ‘k’ means the ‘kth’ set, and ‘L’ means the total number of damaging or LoF heterozygous variants.

### Binomial Analysis

Expected and observed counts of rare variants in each gene were compared by independent binomial tests. The expected number of rare *de novo* damaging variants was determined using the *de novo* expectation model (see above); the expected number of rare transmitted heterozygous variants was determined using the estimation method mentioned above. Inputs for this test were those with inferred pathogenicity, including D-mis (MetaSVM-deleterious or MPC-score ≥ 2) mutations and LOF (stop-gains, stop-losses, frameshift insertions, and deletions, or canonical splice-site mutations) mutations. Binomial analysis for mutational enrichment did not include non-frameshift insertions or deletions, or compound heterozygous variants. The genome-wide significant cutoff was 2.6 × 10^−6^ (= 0.05/19,347).

### Case-control burden analysis

Case and control cohorts were processed using the same pipeline and filtered with the same criteria. A one-sided Fisher’s exact test was used to compare the observed number of total alternative alleles in cases vs. gnomAD controls (without disease-enriched TOPMed samples) for our candidate genes, regardless of transmission pattern.

### Gene lists for specific diseases

Gene lists for VOGM, arteriovenous malformation (AVM), cerebral cavernous malformation (CCM), Moyamoya disease (MMD), and human height were curated from the publications referenced and genes were considered disease-causing based on guidelines from the Centers for Mendelian Genomics ^23, 26, 36, 112, 137–150^. Genes were classified as “possible” VOGM risk genes if they are established VOGM genes, or if they harbored ≥ 1 damaging DNV in a gene with pLI score ≥ 0.9 in the gnomAD database.

### Weighted Gene Co-expression Network Analysis (WGCNA)

Robust consensus WGCNA was conducted by applying a processed bulk transcriptome sequencing data set encompassing sixteen human brain regions across human brain development ^99^. To increase resolution of the primary neurodevelopmental time periods, analysis was limited to timepoints between gestational week 9 and postnatal year 3. We removed samples > 3 standard deviations above the mean network connectivity as outliers. Network analysis was performed with WGCNA ^98^, assigning genes to specific modules based on bi-weight mid-correlations among genes. Soft threshold power of 10 was chosen to achieve scale-free topology (r2 > 0.9). Then a signed co-expression network was generated. The topological overlap matrix was clustered hierarchically using average linkage hierarchical clustering (using ‘1 – TOM‘ as a dis-similarity measure). The topological overlap dendrogram was used to define modules using a minimum module size of 40, a deep split of 4, merge threshold of 0.1.

### Module enrichment analysis

Module gene lists were obtained via WGCNA as described above. In a background set of all genes categorized in co-expression modules, logistic regression was used for an indicator-based enrichment: is.disease ∼ is.module + gene covariates (GC content, gene length, and mean expression in bulk RNA-seq atlas), as described previously ^100^. Of the 88 WGCNA modules, the gray module, by WGCNA convention ^99^, contains all genes that do not co-express and are consequently unassigned to a co-expression network. Thus, the gray module was excluded from enrichment testing, and enrichment significance was defined at the Bonferroni multiple-testing cutoff (α = 5.68 × 10^-04^).

### Cell-type enrichment analysis

Cell-type-enriched genes (cell type markers) were obtained from a scRNA-seq atlas of human brain spanning the period between early fetal development into adulthood and from a scRNA-seq atlas of mouse meninges. In a background set of all genes expressed in ≥3 cells of the scRNA-seq atlas, logistic regression was applied for indicator-based enrichment analysis: is.cell.type ∼ is.disease + gene covariates (GC content, gene length). All p-values were adjusted by Bonferroni correction (α = 1.19 x 10^-03^ for the brain parenchyma and α = 8.33 x 10^-03^ for the meninges).

### Gene Ontology and pathway enrichment analysis

A total of 436 genes including genes harboring damaging DNVs, LoF-intolerant genes (pLI ≥ 0.9) with rare (Bravo MAF ≤ 5 × 10^−5^) damaging transmitted variants, *EPHB4*, *ACVRL1*, and *ENG* were input into EnrichR R package Version 3.0 ^151^. Enrichment analysis was performed for gene ontologies (biological processes, cellular components, and molecular functions), and biological pathway (Wiki pathways and Reactome pathways). The top 10 terms with the lowest adjusted p-values were reported.

### Zebrafish husbandry

Adult *Tg(kdr:gfp)^zn1^* zebrafish (AB background) were maintained in 3.0 L tanks with constant water flow (Iwaki Aquatic) under a 14h/10h light/dark cycle. They were fed GEMMA Micro 500 (Skretting USA) dry food twice daily supplemented with hatched *Artemia* once daily. Embryos were obtained via natural matings; these embryos were maintained at 28.5° C in E3 medium treated with 1-phenyl 2-thiourea (PTU, to prevent melanization for imaging) at a final concentration of 100 µM. Embryos were anesthetized with 0.16 mg/mL tricaine (Syndel). Care protocols were developed from techniques developed at Boston Children’s Hospital ^152, 153^. All animal care and handling was performed in accordance with Yale IACUC protocol 2019-20274.

### CRISPR/Cas9 ribonucleoprotein generation, mRNA synthesis, and injection

Gene-specific crRNA sequences were designed using CRISPOR ^154^ and ordered from IDT. In general, two crRNAs were generated per gene; For paralogous genes such as *ephb4a/b* and *itgb1a/b*, both paralogs were targeted simultaneously. These scRNAs were annealed to tracRNA per the manufacturer’s instructions (to generate complete sgRNAs), then complexed with Cas9 protein at a 1:2 molar ratio (final concentration of RNA:Cas9 was approximately 33 µM:66 µM) in Cas9 buffer to generate active ribonucleoprotein knockout reagents. Phenol red was added to a final concentration of 0.1% w/v to generate injection mix. Embryos at the one-cell stage were injected (WPI Pneumatic PicoPump) with 1 nL of injection mix, then allowed to develop to 48 hpf. In some experiments, these RNPs were coupled with rescue/false-rescue mRNAs. These mRNAs were generated from cloned human cDNAs inserted into pCS2+ and *in vitro* transcribed using mMessage mMachine SP6 kit (ThermoFisher) according to manufacturer’s instructions. These mRNAs were uniformly included in the injection mixes at a concentration of 200 ng/µL.

### Validation of knockout reagents

To verify sgRNA efficacy, injected embryos were collected, and crude genomic DNA was isolated via boiling/alkaline lysis. Primers flanking the sgRNA target site were used to generate an amplicon ∼500 bp in size. This was then denatured, reannealed, and digested with T7 endonuclease (NEB). The resulting digestion product was run out on 2% agarose; sgRNAs were selected fragmentation patterns indicated mismatch DNA arising from CRISPR/Cas9-mediated editing. See **Table S17** for all crRNAs and primer sets used in this study.

### Microangiography, confocal imaging, and statistical analysis

At 48 hpf, zebrafish embryos were anesthetized and injected in the caudal vein with 1 mg/mL 2,000,000 MW rhodamine dextran (ThermoFisher). Embryos were then embedded in 1.5% agarose and immediately imaged on a Leica SP8 confocal microscope. Image stacks were then processed and analyzed using Fiji/ImageJ. All statistical analysis was performed using R. A Welch two-sample t-test was used for all two-level factor analyses; for all multi-level factor analyses, ANOVAs followed by Tukey multiple comparisons of means were employed.

### Plasmids, cell culture, and protein isolation

c-myc-tagged Ephb4 cDNA in pCMV6 was from Origene. Single K650N, R838W and F867L Ephb4 mutations were introduced by site-directed mutagenesis using a QuikChange II XL Site-Directed Mutagenesis Kit (Agilent) according to manufacturer’s instructions. Cos-7 cells (ATCC) were cultured in DMEM supplemented with 10% FBS and 100 U/ml penicillin/streptomycin (all Thermo Fisher Scientific) in 10 cm culture dishes. At 60% confluency, cells were transfected with 10 mg of plasmid using Lipofectamine in Opti-MEM (both Thermo Fisher Scientific). To control for transfection efficiency, cells were co-transfected with 1 µg of pEGFP-N1 plasmid (Takara Bio USA). Where indicated, protein stability was assessed by adding 5 mg/ml cycloheximide (Sigma-Aldrich) to the cell growth medium for the indicated times prior to harvest. Cells were harvested 48 hours after transfection and washed 2 times with ice-cold PBS. GFP content was assessed by flow cytometry using BD Fortessa or BD FACSCanto instruments (BD Biosciences). Proteins were extracted in NP40 Lysis Buffer (NLB, Thermo Fisher Scientific) by 3 repeats of freeze-thawing on dry ice. Extracts were subsequently centrifuged at 13.000 x *g*. for 10 minutes at 4 °C and supernatants were used for immunoprecipitation and western blot analysis.

### Immunoprecipitation and Western blotting

For immunoprecipitation, 20 µg total protein in 200 µl NLB was precleared with 10 µl protein A Dynabeads (Thermo Fisher Scientific) for 20 min at 4 °C. Precleared lysates were incubated with 1 µg goat anti-EPHB4 polyclonal IgG (R&D systems, AF446-SP) at 4°C for 3 hours followed by addition of 10 µl Protein A Dynabeads (Invitrogen) for 20 min to capture the antibody-protein complexes. Beads were washed 5 times for 5 minutes each with 200 µl NLB before protein elution by boiling for 10 minutes in 20 µl 1x NuPage LDS Sample buffer (Thermo Fisher Scientific). For Western blotting, lysates and immunoprecipitates were separated on a Bolt 10% Bis-Tris Plus gel and transferred onto a PVDF membrane (both Thermo Fisher Scientific). To determine total EPHB4 amounts and EPHB4 phosphorylation status the following antibodies were used: goat anti-EPHB4 polyclonal IgG, mouse anti-phosphotyrosine monoclonal IgG (clone 4G10; MilliporeSigma), rabbit anti-ACTB polyclonal IgG (Cell Signaling Technologies, 4967), rabbit anti-TUBB monoclonal IgG (clone 9F3, Cell Signaling Technologies, 2128), and mouse anti-Myc monoclonal IgG (clone 9E10; EMD MilliporeSigma, OP10-200UG), goat anti-rabbit IgG–HRP (Cell Signaling Technology, 7074), horse anti-mouse IgG–HRP (Cell Signaling Technology, 7076) and donkey anti-goat IgG–HRP (Jackson Immunoresearch, 705-035-147).

### CRISPR generation of the EphB4 F867L mouse

Creation of the EphB4 F867L mouse was performed via CRISPR/Cas-mediated genome editing essentially as described ^155, 156^. Potential Cas9 target guide (protospacer) sequences in exon 15 in the vicinity of the F867 codon were screened using the online tool CRISPOR http://crispor.tefor.net, and 5 candidates were selected. Templates for sgRNA synthesis were generated by PCR from a pX330 template, sgRNAs were transcribed *in vitro* (Megashortscript; ThermoFisher) and purified. sgRNA/Cas9 RNPs were complexed and tested for activity by zygote electroporation, incubation of embryos to blastocyst stage, and genotype scoring of indel creation at the target sites. The gRNA with highest activity was selected to create the knock-in allele. A recombination template oligo (IDT) was designed to create the TTT->CTT codon change, and incorporated a P865 CCC->CCT silent mutation to destroy the PAM. sgRNA/Cas9 RNP and the template oligo were electroporated into C57Bl/6J (JAX) zygotes. Embryos were transferred to the oviducts of pseudopregnant CD-1 foster females using standard techniques ^157^. Genotype screening of tissue biopsies from founder pups was performed by PCR amplification and Sanger sequencing, followed by breeding to establish germline transmission of the correctly targeted F867L allele.

Mice carrying the exon 2-exon 3 floxed alleles of Ephb4 have been described ^158^. C*dh5Ert2Cre* mice were obtained from Cancer Research UK ^159^. All mouse experiments complied with Yale and University of Michigan guidelines and were approved by the university IACUC.

### Mouse genotyping

DNA isolated from tail clips was used for genotyping. The following primers were used to show: a) the presence of Cre recombinase: forward 5’-ATTTACTGACCGTACACCAAA-3’ and reverse 5’-CTGTTTTGCACGTTCACCGGC-3’, b) the Ephb4 floxed allele: forward 5′-GGAATGAGGGCGAGTGGGTT-3′ and the reverse 5′-GGTTGGGGACAAAGAGGAAGA-3, and c) the Ephb4 F867L mutant locus: forward 5’-GCGAACGATTCTCTCAAGCC-3’ and reverse 5’-AGAATAGTGAGGCTGCCGTT-3; for the latter, the obtained product was subsequently digested with EcoN1 restriction enzyme (New England Biolabs) to differentiate wild-type, heterozygous and homozygous mice. 20 µl PCR reactions contained 10 µl GoTaq® green PCR Master mix (Promega), 6 µl nuclease-free water, 0.5 µl of each primer (10 µM), and 3 µl template DNA. Amplification was performed as follows: 98°C for 2 minutes, 35 cycles consisting of 98°C for 30 s, 55°C for 30 s, and 72°C for 50 s, followed by 10 min incubation at 72°C. Products were separated on a 2% agarose gel.

### CD31 whole mount staining of yolk sacs and embryos

EPHB4 F867L/+ heterozygous mice were intercrossed, and embryos and yolk sacs were harvested at different stages of embryonic development. Embryos and yolk sacs were fixed in 4x diluted Cytofix, Cytoperm (BD Biosciences) and stained with a rat anti-CD31 antibody (BD Biosciences, 550274), followed by a secondary goat anti-rat Alexa Fluor 488 antibody (Jackson Immunoresearch). Specimens were washed in PBS + 0.3% Triton (Sigma-Aldrich). Images were acquired on a Leica SP5 confocal microscope.

### Embryonic pERK analyses

EPHB4 F867L/+ heterozygous females were crossed with Ephb4^fl/fl^ Cdh5-ert2cre males and pregnant dams were given 2 i.p. injections of tamoxifen (Sigma, 0.05 mg/g body weight, dissolved in corn oil) on consecutive days E13.5 and E14.5 days after the detection of vaginal plugs. Embryos were harvested at E18.5 and whole liver protein extracts were prepared in Cell Lysis Buffer (CLB, Thermo Fisher Scientific). Protein extraction was further facilitated by dounce homogenization, followed by 3 repeats of freeze-thawing on dry-ice. Extracts were subsequently centrifuged at 13,000 x *g* for 10 minutes at 4 °C and pERK levels were determined by Western blotting as above, using rabbit anti-phospho-ERK monoclonal IgG (clone D13.14.E, Cell Signaling Technology, 4370) and rabbit anti-ERK IgG (clone 137F5, Cell Signaling Technology, 4695) and rabbit anti-ACTB polyclonal IgG. Additional embryos were harvested at E18, formalin-fixed, dehydrated and paraffin-embedded. 5 µM sections were stained with a rat anti-CD31 antibody (Dianova, SZ31), followed by secondary goat anti-rat Alexa Fluor 594 (Jackson Immunoresearch). Images were acquired on a Nikon N-SIM + A1R microscope.

### Data and software availability

WES data for all VOGM parent-offspring trios reported in this study have been deposited in the NCBI database of Genotypes and Phenotypes under accession number phs000744.v4.p2.

## Graphical Abstract

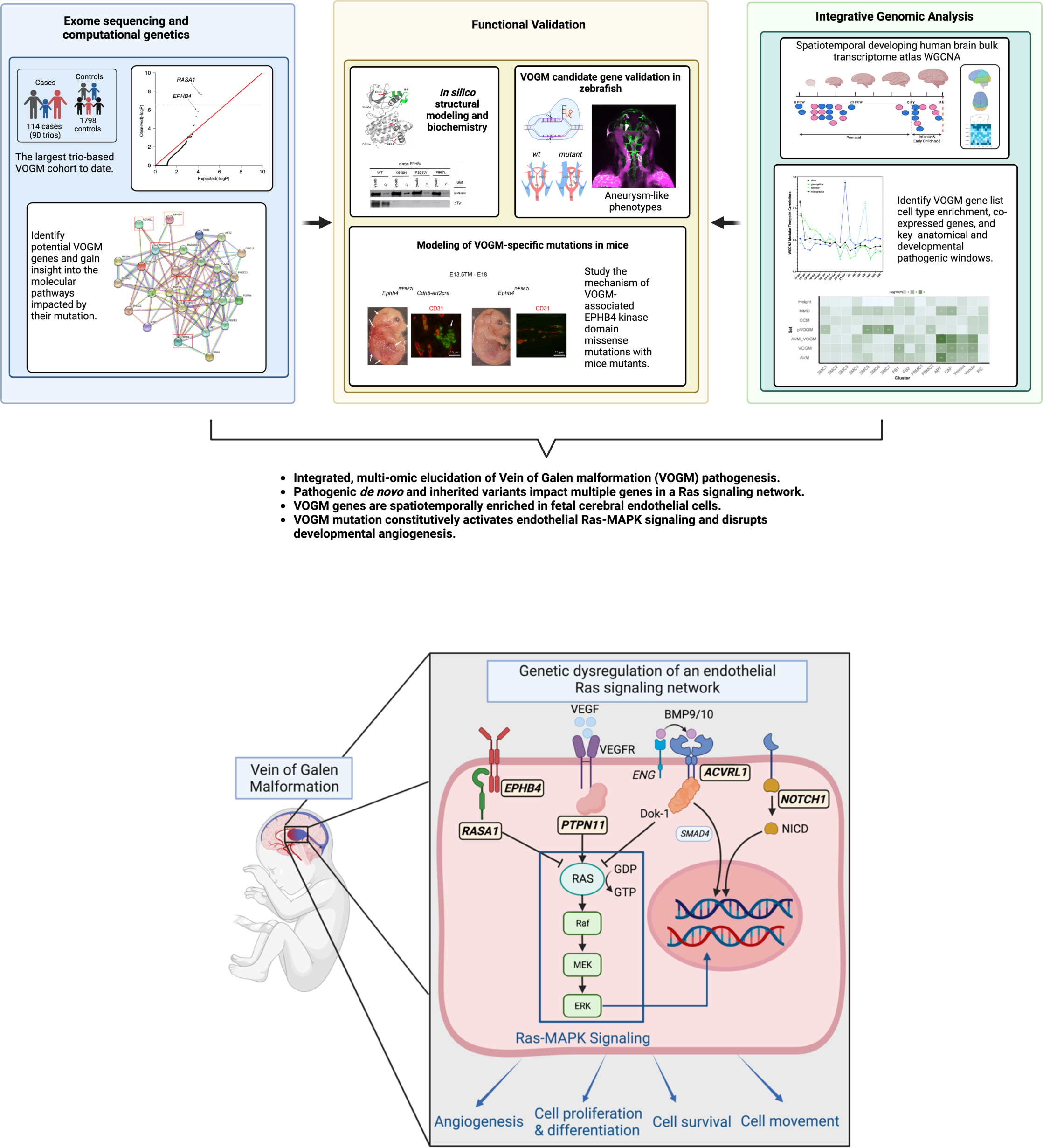

