## Supplementary Figures for "Genetic dysregulation of an endothelial Ras signaling network in vein of Galen malformations"

Brief description of what this file includes:

Supplementary Figure 1: Representative imaging for probands.

Supplementary Figure 2: Mutations in *RASA1* and *EPHB4.*

Supplementary Figure 3: Mutations in *NOTCH1*, *PTPN11* and *ITGB1*.

Supplementary Figure 4: *De novo* mutation rate closely approximates Poisson distribution in VOGM cases and controls.

Supplementary Figure 5: Cutaneous manifestations in VOGM probands and family members.

Supplementary Figure 6: GO term and pathway enrichment analysis.

Supplementary Figure 7: Knockdown of VOGM candidate genes in zebrafish leads to aneurysm-like phenotypes.

Supplementary Figure 8: The remaining top 10 GO biological processes, molecular functions, cellular components, and GO WikiPathways enrichment terms.

Supplementary Figure 9: VOGM gene discovery projections.

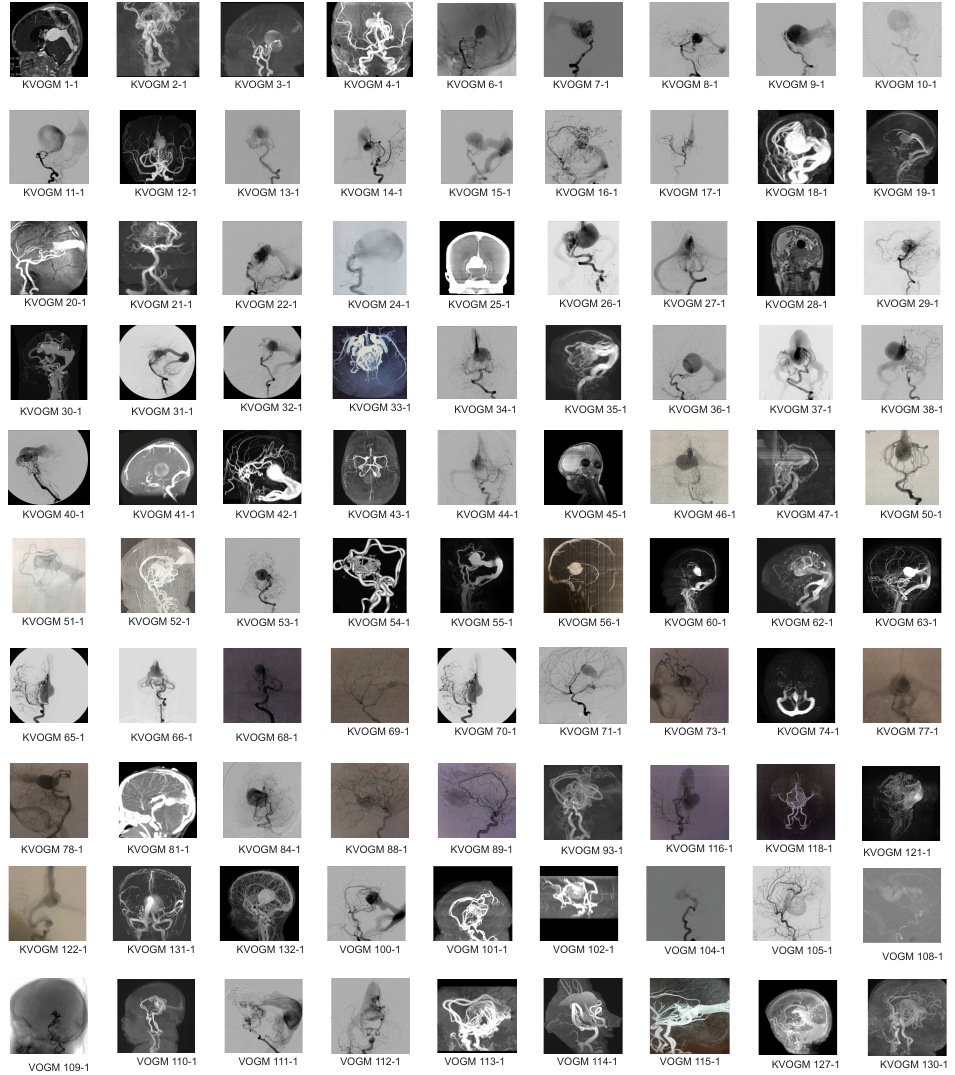

**Supplementary Figure 1. Representative imaging for probands.** Representative images of 3-Tesla time-of-flight magnetic resonance angiography or digital subtraction angiography for all patients with available imaging, with patient codes.

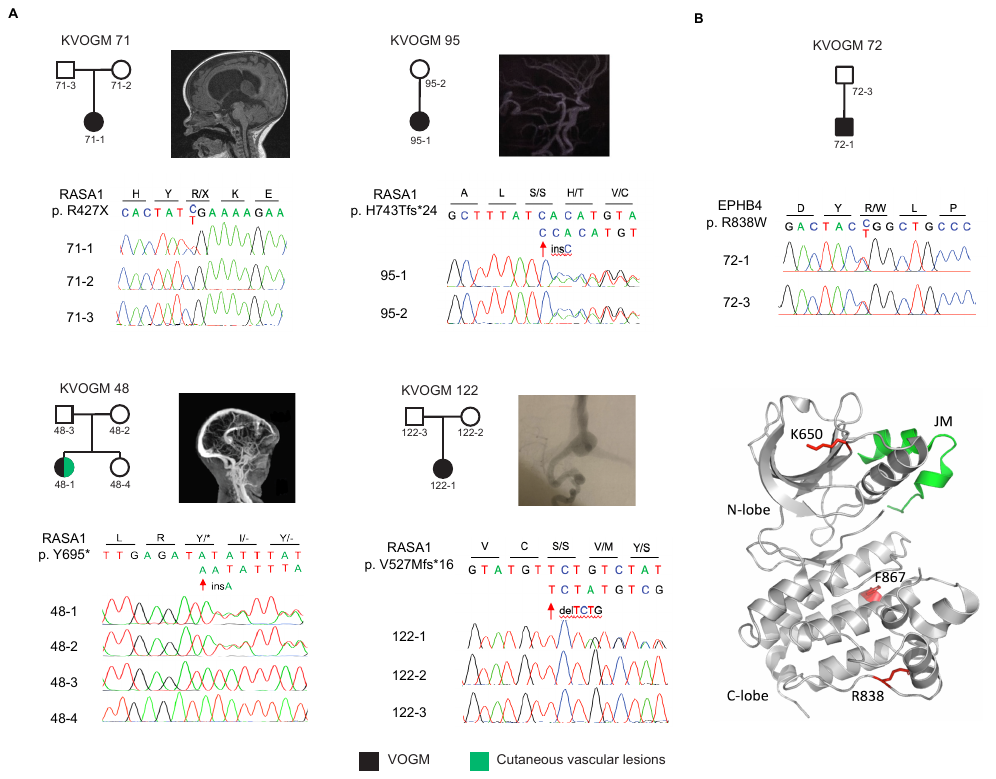

**Supplementary Figure 2. Mutations in *RASA1* and *EPHB4.*** (A) *De novo* and transmitted variants in *RASA1*. Representative digital subtraction angiography reconstructions and pedigrees depicting kindred structure and phenotype are shown. Black symbol represents vein of Galen malformation (VOGM) and green symbols denotes cutaneous vascular lesions. These mutations (p. R427X, p. V527Mfs*16, p. Y695* and p. H743Tfs*24) are confirmed by direct PCR amplification with custom primers followed by Sanger sequencing. For Sanger sequence of variant p. R709X, please refer to our previous publication ^1^. (B)Transmitted variants in *EPHB4*. Representative digital subtraction angiography reconstructions and pedigree depicting kindred structure and phenotype of KVOGM-72 are shown. Variant (p. R838W) been validated by Sanger sequencing. Cartoon diagram of the crystal structure of EphB4 kinase domain (PDB ID: 6FNL) ^2^ in grey. Juxtamembrane region (JM) from an aligned EphB2 structure (PDB ID: 1JPA) ^3^ shown in green. The locations of variants p. K650, p. R838 and p. F867 are indicated in red. For Sanger sequence of variants p. F867L, p. K650N, p. A509G, p. E432Gfs*7, please refer to our previous publication ^1^.

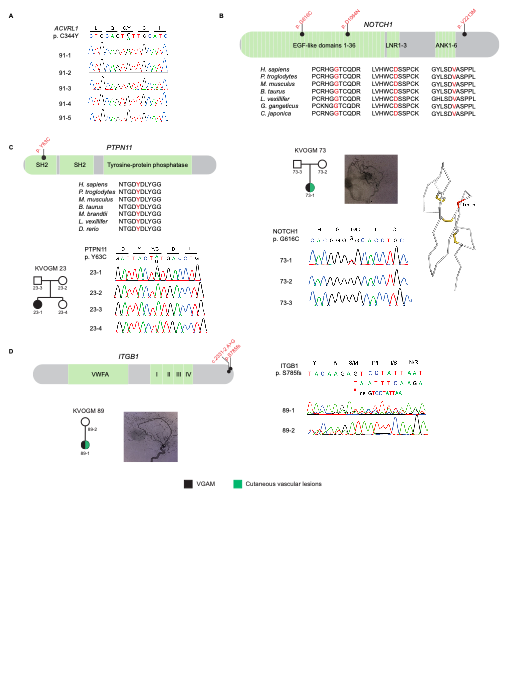

**Supplementary Figure 3. Mutations in *NOTCH1*, *PTPN11* and *ITGB1*.** (A) Multi-generational VOGM family in KVOGM-91. The genotype of the proband and relatives were Sanger-validated. See Figure 2c for pedigree information of this family. (B) Transmitted and de novo variant in *NOTCH1*. Transmitted variant p.V2213M in VGOM46-1, unphased variant p.D1064N in KVOGM83-1, and DNV p.G616C in KVOGM73-1 been mapped to proteins domains. Representative digital subtraction angiography reconstruction and pedigree depicting kindred structure and phenotype for KVOGM73 harboring *de novo* variant G616C. Variant (p. G616C) has been validated by Sanger sequencing. Schematic illustrating the position of G616 within the Alphafold model of this domain (AF-P46531-F1-model_v2.pdb, residues 603-641) of the 16th EGF domain of NOTCH1. Disulfides are indicated and the location of G616 is shown in red. (C) *De novo* variant in *PTPN11*. The variant has been mapped to the SH2 domain. KVOGM23 pedigree depicting kindred structure and phenotype. Gene sequence encoding variant (p. Y63C) has been Sanger-validated. SH2, Src homology 2 domain. (D) Transmitted variants in *ITGB1*. Variants have been mapped to protein domains. Representative digital subtraction angiography reconstruction and pedigree depicting kindred structure and phenotype. Gene sequence encoding variant (p. S785fs) has been Sanger-validated.

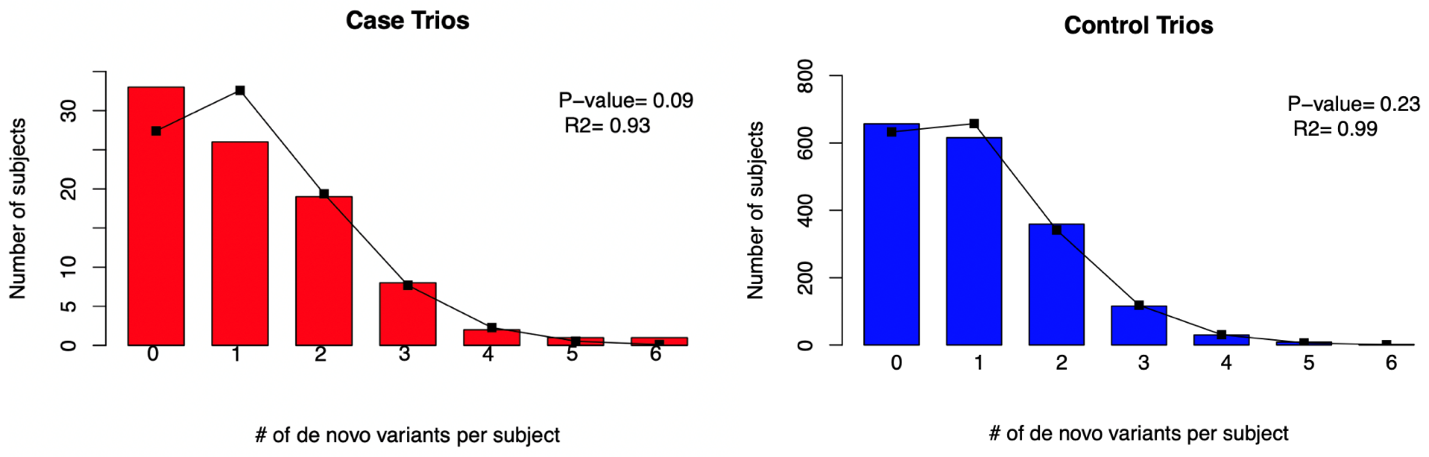

**Supplementary Figure 4*. De novo* mutation rate closely approximates Poisson distribution in VOGM cases and controls.** The observed number of *de novo* mutations per subject (bars) is compared to the numbers expected (line) from the Poisson distribution in the case (red) and control cohorts (blue). ‘p' denotes chi-squared p-value.

**Supplementary Figure 5. Cutaneous manifestations in VOGM probands and family members.** Atypical capillary malformations and other cutaneous vascular lesions are depicted in images of probands and family members, labeled by patient code and familial relation.

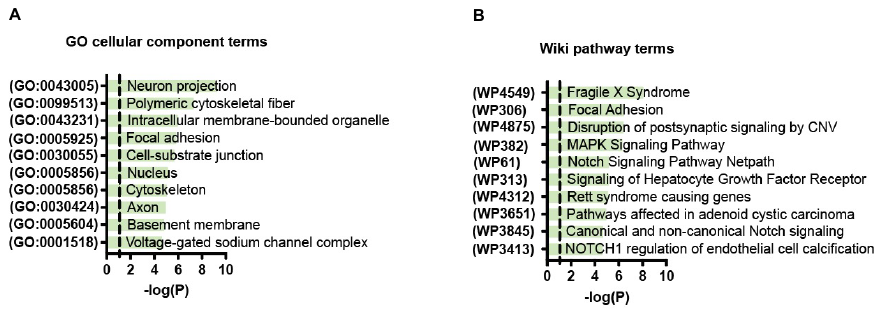

**Supplementary Figure 6. GO term and pathway enrichment analysis.** A) Top 10 GO cellular component and B) GO Wiki pathway enrichment terms. The y-axis depicts GO term or WikiPathways term ID numbers. The x axis depicts -log (P value) and the dotted line represents the α = 0.05 significance threshold. The GO term and WikiPathways term name overline their respective bars.

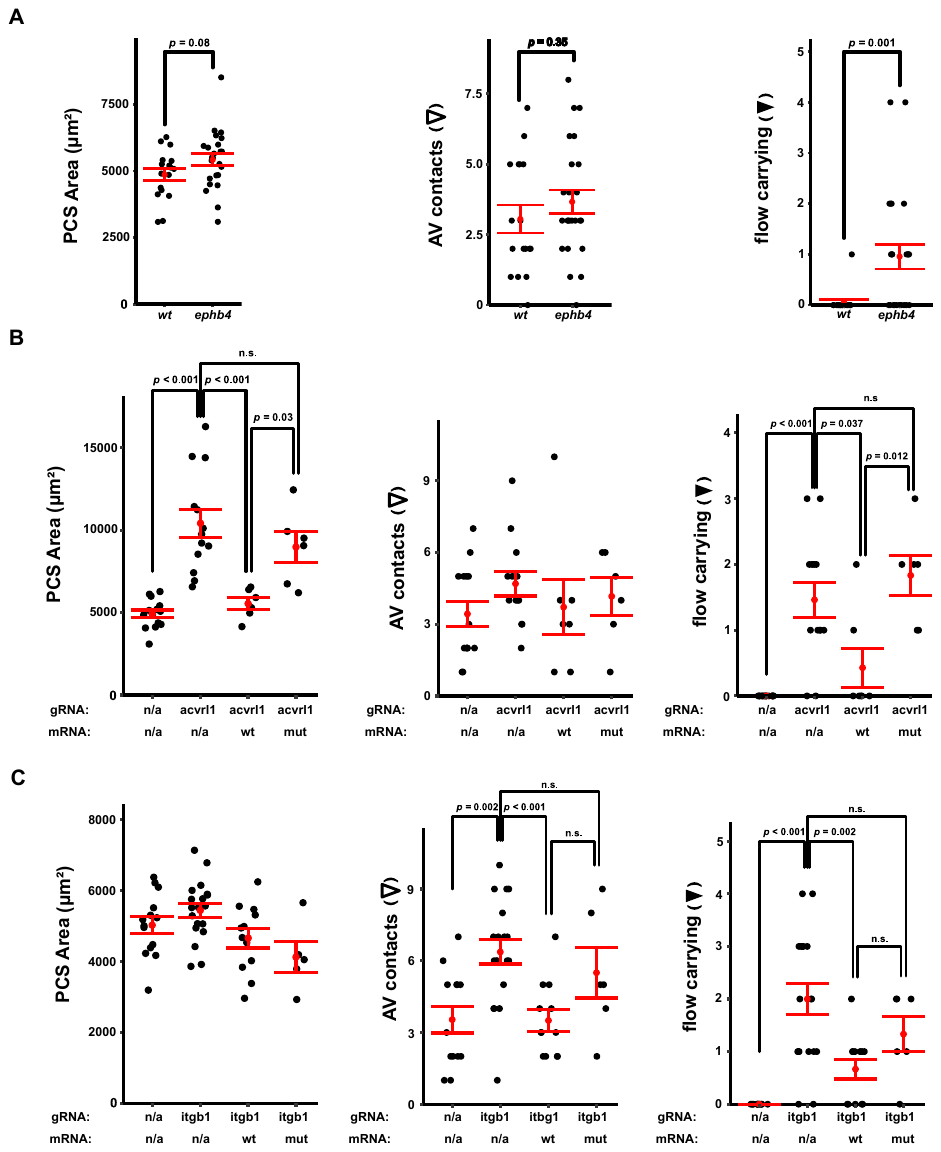

**Supplementary Figure 7. Knockdown of VOGM candidate genes in zebrafish leads to aneurysm-like phenotypes.** (A) Quantifications of the PCS area, AV contacts, and flow-carrying AVs in *ephb4a/b* loss-of-function experiments for 48 hpf wt and ephb4a/b-depleted zebrafish embryos. Loss of *ephb4a/b* function does not significantly increase PCS size (p = 0.08), does not significantly increase AV count (p = 0.35), but significantly increases flow-carrying AVs (p = 0.001). (B) Quantifications for the *acvrl1* loss-of-function experiments of 48 hpf wt, *acvrl1*-depleted, rescue (with acvrl1+ mRNA) and false rescure (with acvrl1 mutant mRNA) zebrafish embryos. Loss of *acvrl1* function significantly increases PCS size (p < 0.001) and is rescued by *wt* *acvrl1* mRNA (p < 0.001) but not by mutant *acvrl1* mRNA (p = 0.03). Quantification of AVs in *acvrl1* loss-of-function/rescue experiments showing loss of *acvrl1* function does not affect the number of AV contacts across all conditions. Loss of *acvrl1* function significantly increases flow-carrying AV count (p < 0.001). This can be rescued by *wt acvrl1* (p = 0.037) but not mutant *acvrl1* mRNA (n.s. vs. loss-of-function condition). (C) Quantifications for the *ITGB1* loss-of-function experiments of 48 hpf wt *itgb1*-depleted, rescue (with *ITGB1*+ mRNA) and false rescue (with *ITGB1* mutant mRNA) zebrafish embryo. Loss of *itgb1a/b* function does not affect PCS area across all conditions, but significantly increases AV contacts (p = 0.002) shown in ITGB1 mutant. This increase is rescued by *wt itgb1a/b* (p < 0.001), but not by mutant *itgb1a/b* mRNA (not significant relative to loss-of-function condition). Loss of *itgb1a/b* function significantly increases flow-carrying AV count (p < 0.001) and is rescued by *wt itgb1a/b* (p = 0.002), but not by mutant *itgb1a/b* mRNA (n.s. vs. loss-of-function condition).

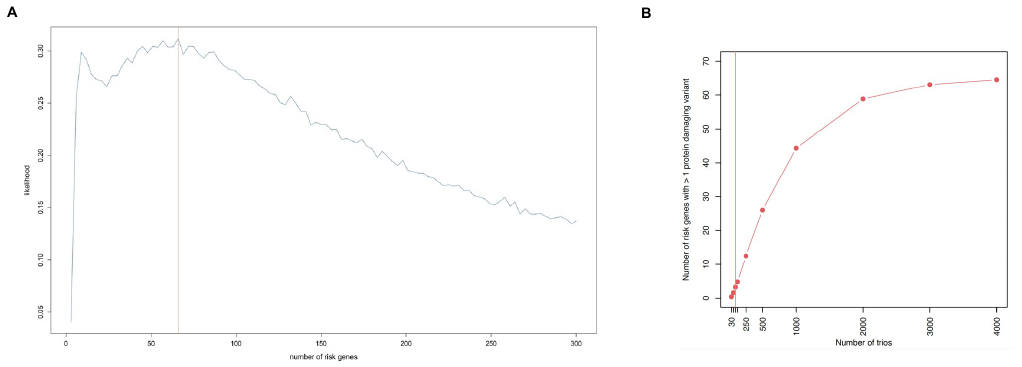

**Supplementary Figure 9. VOGM gene discovery projections.** (A) Estimated number of VOGM risk genes via a *de novo* mechanism. A Monte Carlo simulation was performed based on observed protein-altering *de novo* variants in 3,049 loss-of-function intolerant genes (pLI ≥ 0.9 in gnomAD [v2.1.1]) using 20,000 iterations. We estimate the number of risk genes via a *de novo* mechanism is ~66. (B) Estimated number of genes with more than one protein-damaging variant with an increasing number of trios. The number of trios is specified on the x-axis and the number of genes with more than one protein-damaging *de novo* variant on the y-axis. The expected rate of protein-altering *de novo* variants was modeled in 10,000 iterations with an increasing number of trios, given the probability of *de novo* protein-altering variants. WES of 250 and 1,000 trios are expected to yield a saturation rate of 20.1% and 70.5% respectively, for all VOGM risk genes.
